# Individuality transfer: Predicting human decision-making across task conditions

**DOI:** 10.1101/2025.03.25.645375

**Authors:** Hiroshi Higashi

## Abstract

Predicting an individual’s behaviour in one task condition based on their behaviour in a different condition is a key challenge in modeling individual decision-making tendencies. We propose a novel framework that addresses this challenge by leveraging neural networks and introducing a concept we term the “individual latent representation.” This representation, extracted from behaviour in a “source” task condition via an encoder network, captures an individual’s unique decision-making tendencies. A decoder network then utilizes this representation to generate the weights of a task-specific neural network (a “task solver”), which predicts the individual’s behaviour in a “target” task condition. We demonstrate the effectiveness of our approach in two distinct decision-making tasks: a value-guided task and a perceptual task. Our framework offers a robust and generalizable approach for parameterizing individual variability, providing a promising pathway toward computational modeling at the individual level—replicating individuals in silico.

## 1 Introduction

Humans (and other animals) exhibit substantial commonalities in their decision-making processes. However, considerable variability is also frequently observed in how individuals perform perceptual and cognitive decision-making tasks [5, 1]. This variability arises from differences in underlying cognitive mechanisms. For example, individuals may vary in their ability or tendency to retain past experiences [12, 8], respond to events with both speed and accuracy [52, 45], or explore novel actions [17]. If these factors can be meaningfully disentangled, they would enable a concise characterization of individual decision-making processes, yielding a low-dimensional, parameterized representation of individuality. Such a representation could, in turn, be leveraged to predict future behaviours at an individual level. Shifting from population-level predictions to an individual-based approach would mark a significant advancement in domains where precise behaviour prediction is essential, such as social and cognitive sciences. Beyond prediction, this approach offers a framework for parameterizing and clustering individuals, thereby facilitating the visualization of behavioural heterogeneity, which has applications in psychiatric analysis [31, 10]. Furthermore, this parameterization offers a promising pathway toward computational modeling at the individual level—replicating the cognitive and functional characteristics of individuals *in silico* [41].

Cognitive modelling is a standard approach for reproducing and predicting human behaviour [29, 3, 57], often implemented within a reinforcement learning framework (e.g., [30, 9, 54]). However, because these cognitive models are manually designed by researchers, their ability to accurately fit behavioural data may be limited [16, 44, 28, 13]. A data-driven approach using artificial neural networks (ANNs) offers an alternative [11, 33, 39]. Unlike cognitive models, which rely on predefined behavioural assumptions [37], ANNs require minimal prior assumptions and can learn complex patterns directly from data. For instance, convolutional neural networks (CNNs) have successfully replicated human choices and reaction times in various visual tasks [25, 35, 15]. Similarly, recurrent neural networks (RNNs) [42, 7] have been applied to model value-guided decision-making tasks such as the multi-armed bandit problem [56, 10]. A promising approach to capturing individual decision-making tendencies while preserving behavioural consistency is to tune ANN weights using a parameterized representation of individuality.

This idea was first proposed by Dezfouli et al. [10], who employed an RNN to solve a two-armed bandit task. Their study utilized an autoencoder framework [38, 48], in which behavioural recordings from a single session of the bandit task, performed by an individual, were fed into an encoder. The encoder produced a low-dimensional vector, interpreted as a latent representation of the individual. Similar to hypernetworks [20, 22], a decoder then took this low-dimensional vector as input and generated the weights of the RNN. This framework successfully reproduced behavioural recordings from other sessions of the same bandit task while preserving individual characteristics. However, since this individuality transfer has only been validated within the bandit task, it remains unclear whether the extracted latent representation captures an individual’s intrinsic tendencies across a variety of task conditions.

To address this question, we aim to make the low-dimensional representation— referred to as the *individual latent representation*—robust to variations across individuals and task conditions, thereby enhancing its generalizability. Specifically, we propose a framework that predicts an individual’s behaviours, not only in the same condition but also in similar yet distinct task conditions and environments. If the individual latent representation serves as a low-dimensional representation of an individual’s decision-making process, then extracting it from one condition could facilitate the prediction of that individual’s behaviours in another.

In this study, we define the problem of *individuality transfer across task conditions* as follows (also illustrated in Figure 1). We assume access to a behavioural dataset from multiple individuals performing two task conditions: a

**Figure 1.**
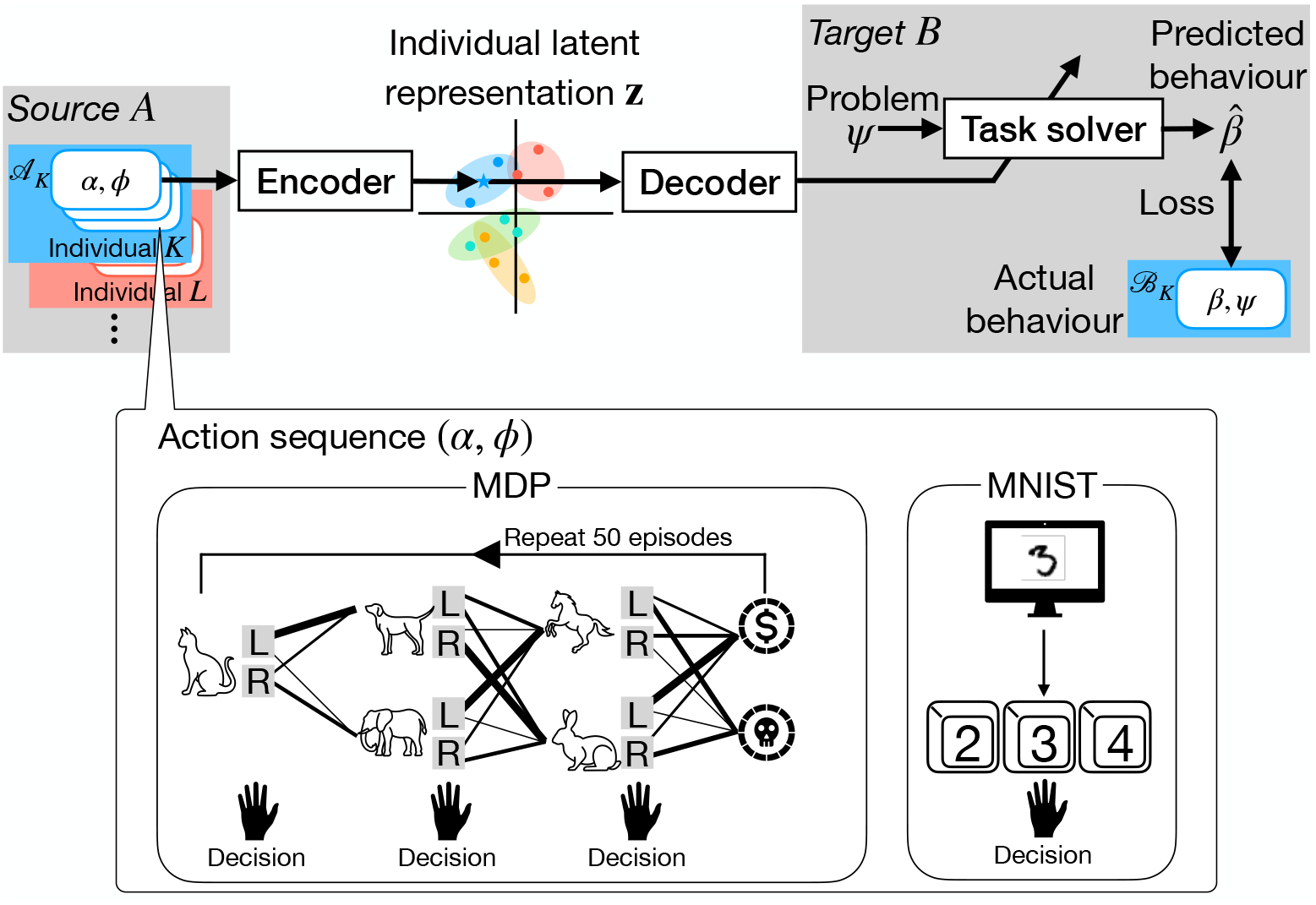
The EIDT (encoder, individual latent representation, decoder, and task solver) framework for individuality transfer across task conditions. The encoder maps action(s) *α*, provided by an individual *K* performing a specific problem *ϕ* in the source task condition *A*, into an individual latent representation (represented as a point in the two-dimensional space in the center). The individual latent representation is then fed into the decoder, which generates the weights for a task solver. The task solver predicts the behaviour of the same individual *K* in the target task condition *B*. During the training, a loss function evaluates the discrepancy between the predicted behaviour 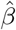 and the actual recorded behaviour *β* of individual *K*. The encoder’s input is referred to as an *action sequence*, the form of which depends on task. For example, in a sequential Markov decision process (MDP) task, an action sequence consists of an environment (state transition probabilities) and a sequence of actions over multiple episodes. For a digit recognition task, it consists of a stimulus digit image and the corresponding chosen response.

### *source task condition* and a *target task condition*

We train an encoder that takes behavioural data from the source task condition as input and outputs an individual latent representation. This representation is then fed into a decoder, which generates the weights of an ANN, referred to as a *task solver*, that reproduces behaviours in the target task condition. For testing, a new individual provides behavioural data from the source task condition, allowing us to infer his/her individual latent representation. Using this representation, a task solver is constructed to predict how the test individual will behave in the target task condition. Importantly, this prediction does not require any behavioural data from the test individual performing the target task condition. We refer to this framework as *EIDT*, an acronym for encoder, individual latent representation, decoder and task solver.

We evaluated whether the proposed EIDT framework can effectively transfer individuality in both value-guided sequential decision-making tasks and perceptual decision-making tasks. To assess its generalizability across individuals, meaning its ability to predict the behaviour of previously unseen individuals, we tested the framework using a test participant pool that was not included in the dataset used for model training. To determine how well our framework captures each individual’s unique behavioural patterns, we compared the prediction performance of a task solver specifically designed for a given individual with the performance of task solvers designed for other individuals. Our results indicate that the proposed framework successfully mimics decision-making while accounting for individual differences.

## 2 Results

We evaluated our EIDT framework using two distinct experimental paradigms: a value-guided sequential decision-making task (MDP task) and a perceptual decision-making task (MNIST task). For each paradigm, we assessed model performance in two scenarios. The first, *Within-Condition Prediction*, tested a model’s ability to predict behaviour within a single task condition without individuality transfer. In this scenario, a model was trained on data from a pool of participants to predict the behaviour of a held-out individual in that same condition. The second, *Cross-Condition Transfer*, tested the core hypothesis of individuality transfer. Here, a model used behavioural data from a participant in “source” condition to predict that same participant’s behaviour in a different “target” condition.

The prediction performance was evaluated using two metrics: the negative log-likelihood on a trial-by-trial basis, and the rate for behaviour matched. The negative log-likelihood is based on the probability the model assigned to the specific action that the human participant actually took on that trial. The rate for behaviour matched measures the proportion of trials where the model’s most likely action (deterministically predicted by sampling from the output probabilities) matched the participant’s actual choice.

### 2.1 Markov decision process (MDP) task

The dataset consisted of behavioural data from 81 participants who performed both 2-step and 3-step MDP tasks. Each participant completed three blocks of 50 episodes for each condition, resulting in 486 action sequences in total. All analyses were performed using a leave-one-participant-out cross-validation procedure. For each fold, the model was trained on 80 participants, with 90% used for training updates and 10% for validation-based early stopping.

#### Task solver accurately predicts average behaviour

First, we validated our core neural network architecture in Within-Condition Prediction. We trained a standard task solver, using the architecture defined in Section 4.2.4, on the training/validation pool (*N* = 80) to predict the behaviour of the held-out participant. We compared its performance against a standard cognitive model (a Q-leaning model, Section 4.2.3) whose parameters were averaged from fits to the same training/validation pool.

As shown in Figure 2, the neural network-based task solver significantly outperformed the cognitive model. A two-way (model: cognitive model/task solver, task condition: 2-step/3-step) repeated-measures (RM) ANOVA with Greenhouse-Geisser correction (significant level was 0.05) revealed a significant effect of the model on both negative log-likelihood (model: *F*_1,80_ = 148.828, *p* < 0.001, 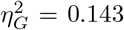, task condition: *F*_1,80_ = 1.107, *p* = 0.296, 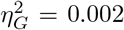, interaction: *F*_1,80_ = 0.240, *p* = 0.626, 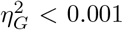 and the rate for behaviour matched (model: *F*_1,80_ = 110.684, 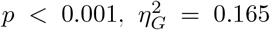, task condition: *F*_1,80_ = 3.914, *p* = 0.051, 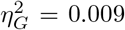, interaction: *F*_1,80_ = 19.059, *p* < 0.001, 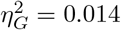). This result confirms that our RNN-based architecture serves as a strong foundation for modeling decision-making in this task.

**Figure 2.**
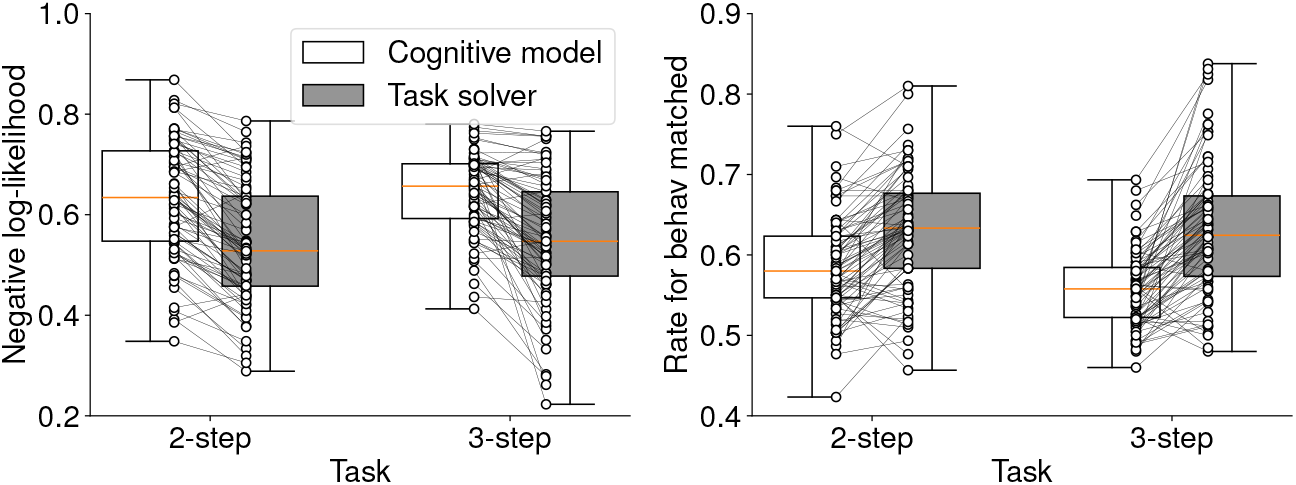
Comparison of prediction performance in Within-Condition Prediction for the MDP task. The plots show the negative log-likelihood (left) and the rate for behaviour matched (right) for the average-participant cognitive model and the task solver for 2-step and 3-step conditions. Box plots indicate the median and interquartile range. Whiskers extend to the minimum and maximum values. Each connected pair of dots represents a single participant’s data. The task solver demonstrates significantly better performance.

#### EIDT enables accurate individuality transfer

Next, we tested our main hypothesis in Cross-Condition Transfer. We used the full EIDT framework to predict a participant’s behaviour in a target condition (e.g., 3-step MDP) using their behavioural data from a source condition (e.g., 2-step MDP). We compared the performance of two models:

#### Cognitive mode

A Q-learning model whose parameters (*q*_lr_, *q*_init_, *q*_dr_, and *q*_it_) were individually fitted for each participant using their data from the source condition and then applied to predict behaviour in the target condition.

#### EIDT

Our framework, trained on the training and validation pool using data from both source and target conditions (see Figure S2, Supplementary Materials for representative training and validation curves). To predict behaviour for a test participant, their individual latent representation was computed by averaging the encoder’s output across all of their behavioural sequences from the source condition, and this representation was fed to the decoder to generate the task solver weights. For reference, the averaged individual latent representations are visualized in Figure S3, Supplementary Materials.

The EIDT framework demonstrated significantly better prediction accuracy than the individualized cognitive model (Figure 3). A two-way (model: cognitive model/EIDT, transfer direction: 2→3/3→2) RM ANOVA confirmed a significant effect of the model on negative log-likelihood (model: *F*_1,80_ = 95.705, *p* < 0.001, 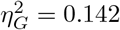, transfer direction: *F*_1,80_ = 14.255, 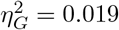, interaction: *F*_1,80_ = 0.008, *p* = 0.012, 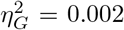) and the rate for behaviour matched (model: *F*_1,80_ = 100.843, *p* < 0.001, 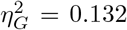, transfer direction: *F*_1,80_ = 13.021, *p* = 0.001, 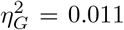, interaction: *F*_1,80_ = 0.964, *p* = 0.329, 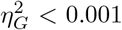). This result indicates that EIDT successfully captures and transfers individual-specific behavioural patterns more effectively than a traditional parameter-based transfer approach.

**Figure 3.**
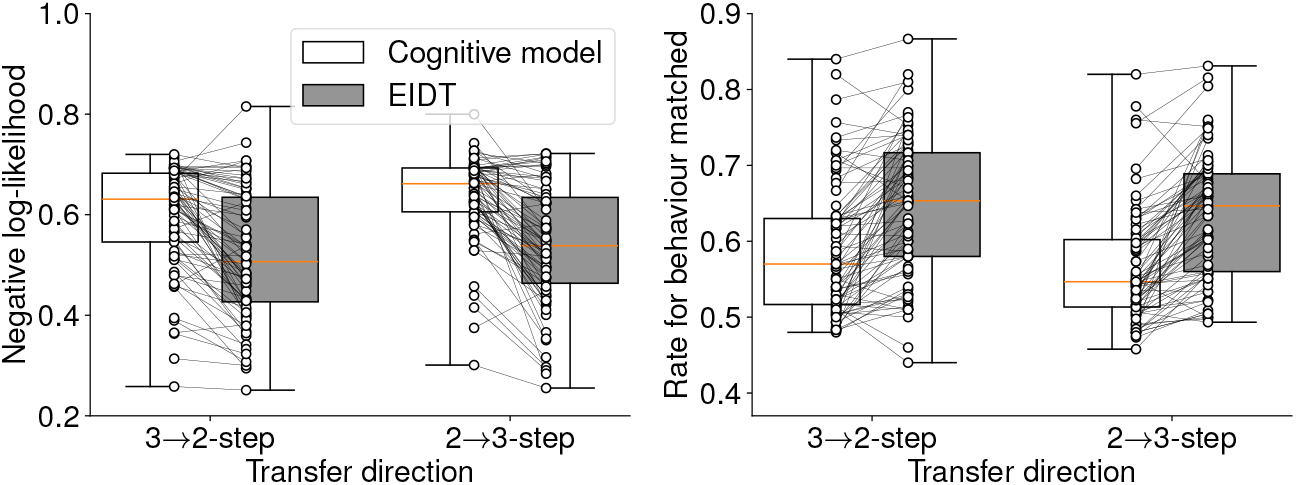
Individuality transfer performance in Cross-Condition Transfer for the MDP task. The plots compare the EIDT framework against an individualized cognitive model on negative log-likelihood (left) and rate for behaviour matched (right) for both 2-step to 3-step and 3-step to 2-step transfer. Box plots indicate the median and interquartile range. Whiskers extend to the minimum and maximum values. Each connected pair of dots represents a single participant’s data. The EIDT model shows superior prediction accuracy.

#### Latent space distance predicts transfer performance

To verify that the individual latent representation meaningfully captures individuality, we conducted a “cross-individual” analysis. We generated a task solver using the latent representation of one participant (Participant *l*) and used it to predict the behaviour of another participant (Participant *k*). We then measured the relationship between the prediction performance (*y*_*k,l*_) and the Euclidean distance (*d*_*k,l*_) between the latent representations of Participants *k* and *l*. As hypothesized, prediction performance was strongly dependent on this distance (Figure 4). We fitted the data using a generalized linear model (GLM): *y*_*k,l*_ ~ Gamma log(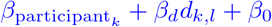). The fitting confirmed that distance (*d*_*k,l*_) was a significant predictor: the coefficient *β*_*d*_ was significantly positive for negative log-likelihood (transfer direction 3→2: *β*_*d*_ = 0.176, *p* < 0.001, 2→3: *β*_*d*_ = 0.316, *p* < 0.001) and significantly negative for the rate for behaviour matched (3 → 2: *β*_*d*_ = 0.106, *p* < 0.001, 2 →3: *β*_*d*_ = − 0.149, *p* < 0.001). This indicates that prediction performance degrades as the behavioural dissimilarity (represented by distance in the latent space) between the source and target individual increases, providing direct evidence that the latent space organizes individuals by behavioural similarity.

**Figure 4.**
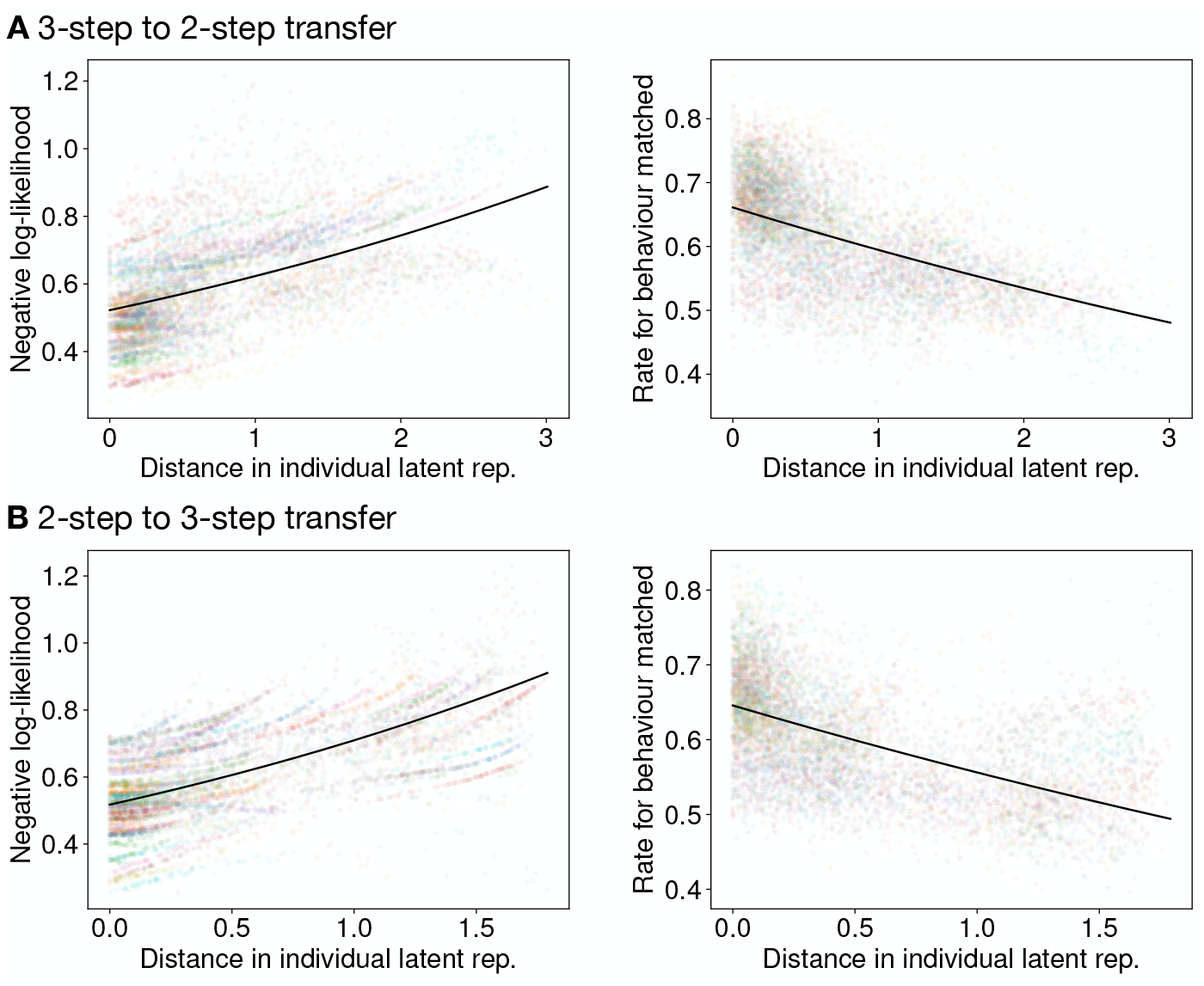
Prediction performances as a functions of latent space distance in the MDP task. This cross-individual analysis shows the result of using a task solver generated from one participant to predict the behaviour of another participant. The horizontal axis is the Euclidean distance between the latent representation of the two participants. The vertical axis shows the negative log-likelihood (left) and rate for behaviour matched (right). Each dot represents one participant pair. Performance degrades as the distance between individuals increases, with the solid line showing the GLM fit. **A** 3-step to 2-step transfer. **B** 2-step to 3-step transfer.

#### On-policy simulations generate human-like behaviour

To assess if our model could generate realistic behaviour, we conducted on-policy simulations. Task solvers specialized to each individual via EIDT performed the MDP task using the same environments as the human participants. We compared the model behaviour to human behaviour on two metrics: total reward per block and the rate of highly-rewarding action selected in the final step.

The model-generated behaviours closely mirrored human behaviours (Figure 5). We found significant correlations between humans and their corresponding models in both total rewards (3 → 2: *R* = 0.667, *p* < 0.001; 2 → 3: *R* = 0.593, *p* < 0.001) and the rate of highly-rewarding action selected (3 → 2: *R* = 0.889, *p* < 0.001; 2 → 3: *R* = 0.835, *p* < 0.001). This demonstrates that the EIDT framework captures individual tendencies that generalize to active, sequential behaviour generation.

**Figure 5.**
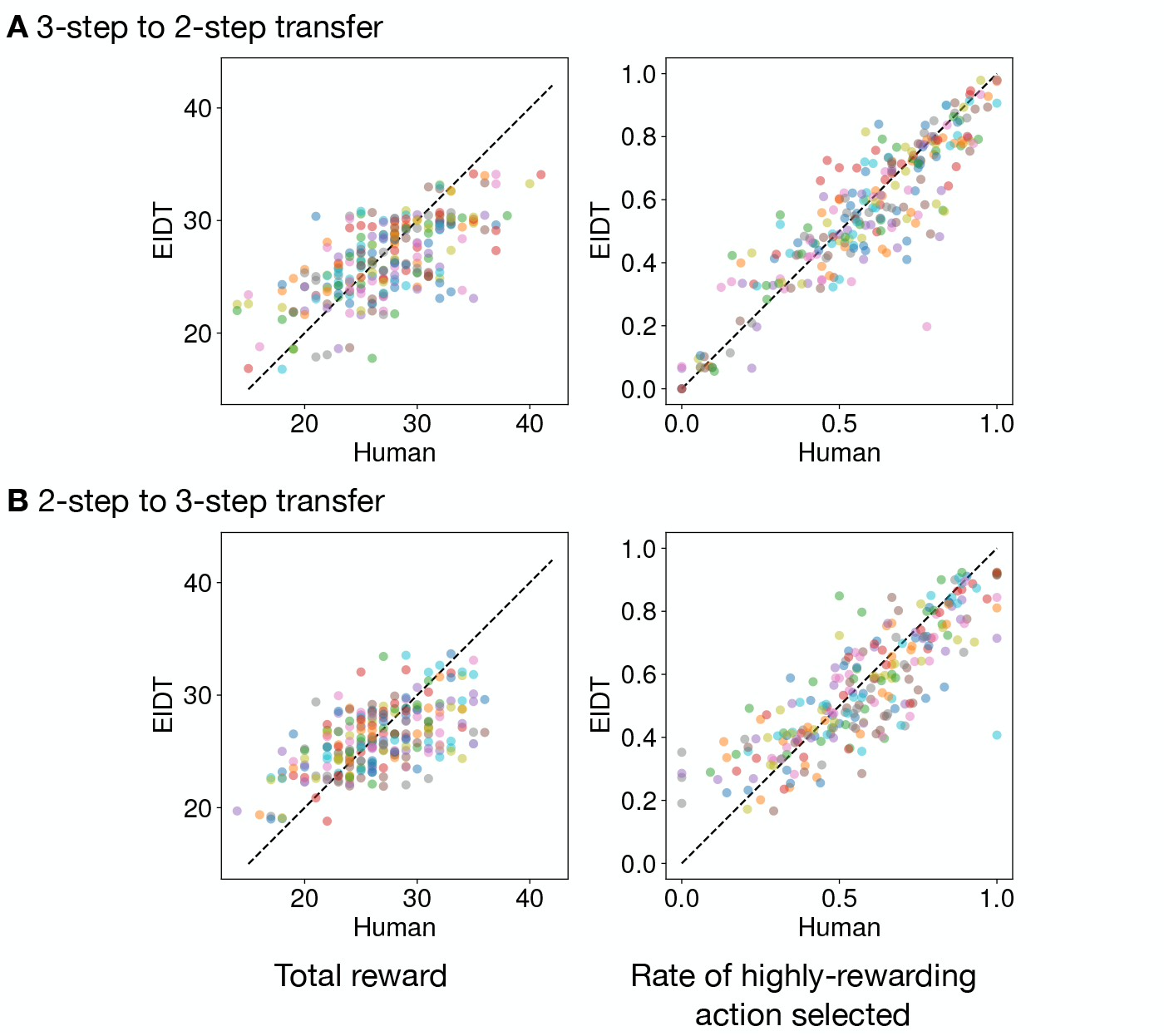
Comparison of on-policy behaviour between humans and EIDT-generated task solvers. Each dot represents the performance of a single human participant (horizontal axis) versus their corresponding model (vertical axis) for one block. Plots show the total reward (left) and the rate of highly-rewarding action selected (right). **A** 3-step to 2-step transfer. **B** 2-step to 3-step transfer.

#### Individual latent representations reflect cognitive parameters

To better interpret the latent space, we applied our EIDT model (trained only on human data) to simulated data from 1,000 Q-learning agents. The agents had known learning rates (*q*_lr_) and inverse temperatures (*q*_it_) sampled from distributions matched to human fits (Figure S1, Supplementary Materials). A cross-individual analysis on these agents confirmed that latent space distance predicted performance, mirroring the results from human data (Figure S5, Supplementary Materials).

The results revealed a systematic mapping between the cognitive parameters and the coordinates of the individual latent representation (Figures 6 and S4, Supplementary Materials). A GLM analysis (Table S1, Supplementary Materials) showed that both both *q*_rl_ and *q*_it_ (and their interaction) were significant predictors of the latent dimensions (*z*_1_ and *z*_2_). This indicates that our datadriven representation captures core computational properties defined in classic reinforcement learning theory.

**Figure 6.**
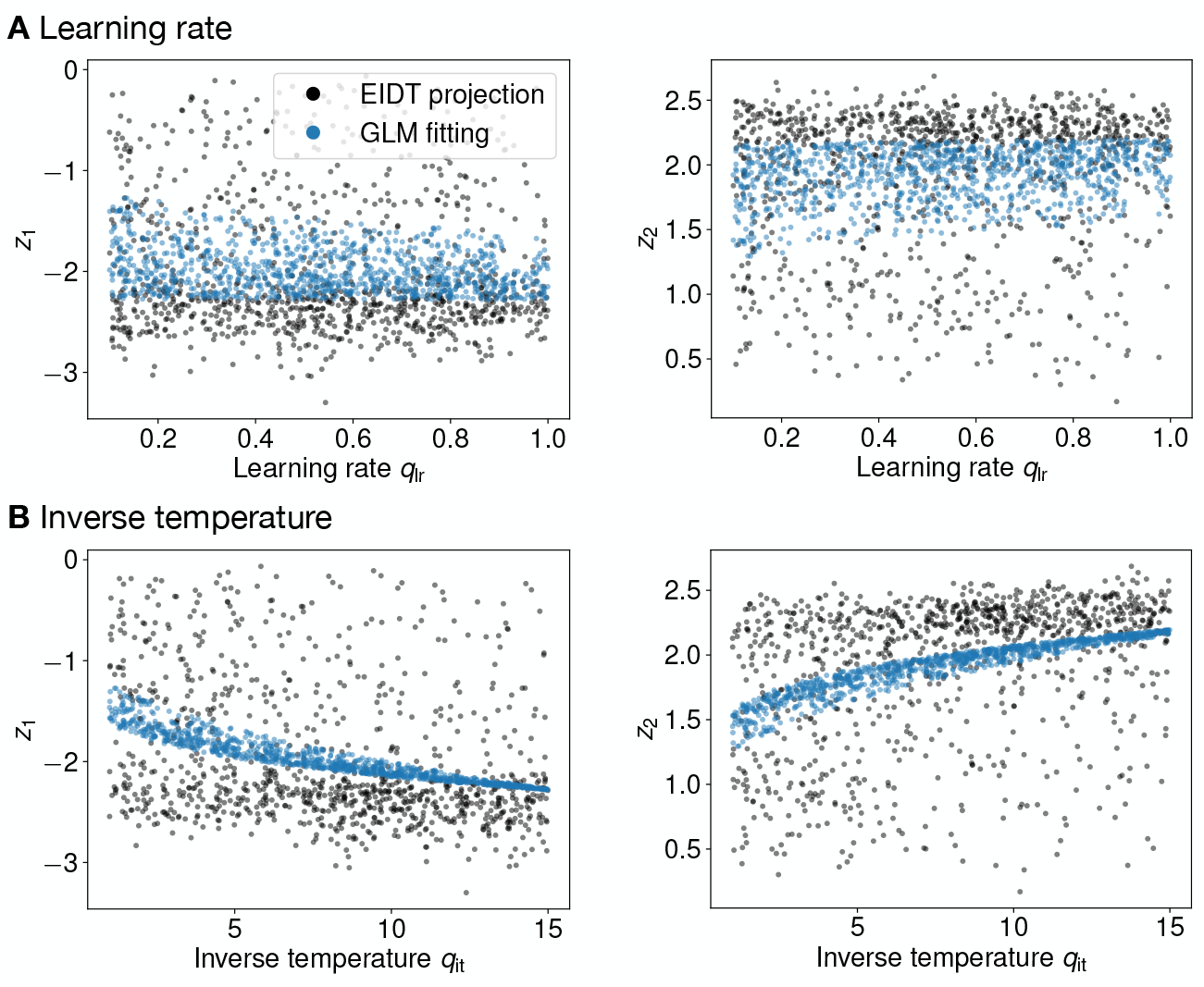
Mapping of Q-learning parameters to the individual latent space for the 3-step MDP task. Each plot shows one dimension of the latent representation (*z*_1_ (left) or *z*_2_ (right)) as a function of either the learning rate (*q*_lr_, **A**) or the inverse temperature (*q*_it_, **B**) of simulated Q-learning agents. Black dots represent the latent representation produced by the encoder from the agent’s behaviour. Blue dots show the fit from a GLM.

### 2.2 Handwritten digit recognition (MNIST) task

We then sought to replicate our findings in a different domain: perceptual decision-making. We used data from Rafiei et al. [34], where 60 participants identified noisy images of digits under four conditions varying in difficulty and speed-accuracy focus (EA: easy, accuracy focus, ES: easy, speed focus, DA: difficult, accuracy focus, and DS: difficult, speed focus). Analyses were again conducted using leave-one-participant-out cross-validation.

#### Task solver outperforms RTNet

First, in Within-Condition Prediction, our base task solver demonstrated task performance (rate of correct responses indicating how accurately a human participant or model responded to the stimulus digit) comparable to human participants and established RTNet model [34] (Figure 7). A two-way (model: human/RTNet/Task solver, task condition: EA/ES/DA/DS) RM ANOVA showed no significant effect of model type (*F*_2,118_ = 1.546, *p* = 0.219, 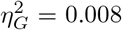), while the task condition had a significant effect (*F*_3,177_ = 866.322, *p* < 0.001, 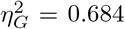). This confirms similar task-solving ability.

**Figure 7.**
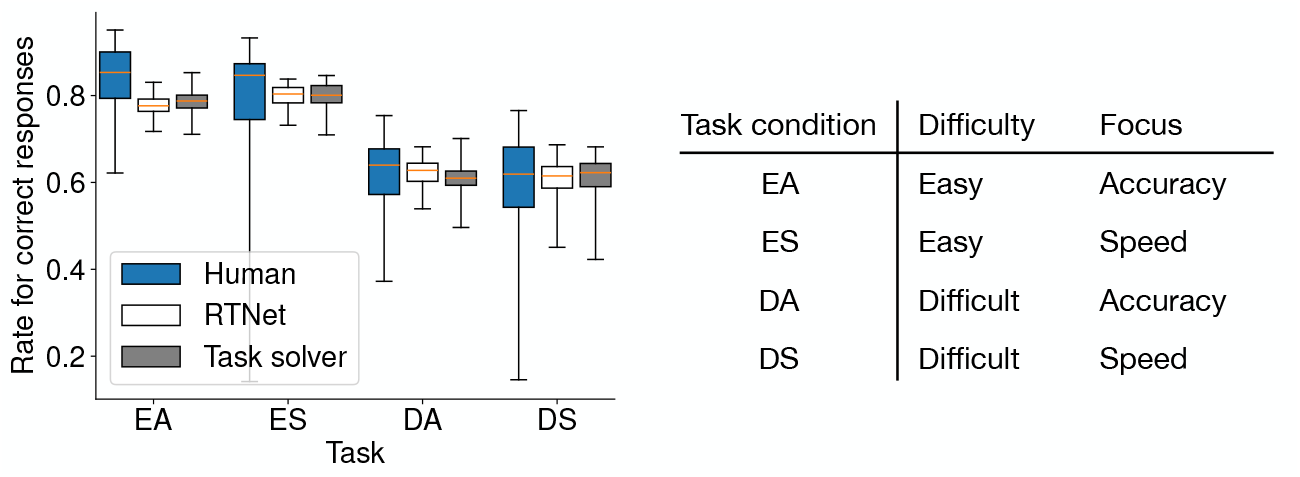
Task performance (rate of correct responses) in Within-Condition Prediction for the MNIST tasks. Box plots indicate the median and interquartile range. Whiskers extend to the minimum and maximum values. Performance is compared across human participants, the RTNet model, and our task solver for the four experimental conditions (EA, ES, DA, and DS). All three show similar performance patterns.

However, the task solver significantly outperformed RTNet in predicting participants’ trial-by-trial choices (Figure 8). A two-way RM ANOVA revealed significant effects of on both negative log-likelihood (model: *F*_1,59_ = 1312.328, *p* < 0.001, 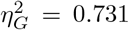, task condition: *F*_3,177_ = 460.535, *p* < 0.001, 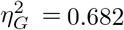, their interaction: *F*_3,177_ = 24.476, *p* < 0.001, 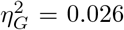) and the rate for behaviour matched (model: *F*_1,59_ = 43.544, *p* < 0.001, 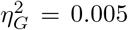, task condition: *F*_3,177_ = 455.728, *p* < 0.001, 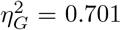, their interaction: *F*_3,177_ = 11.052, *p* < 0.001, 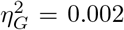). This confirms the task solver’s suitability for modeling individual behaviour in this task.

**Figure 8.**
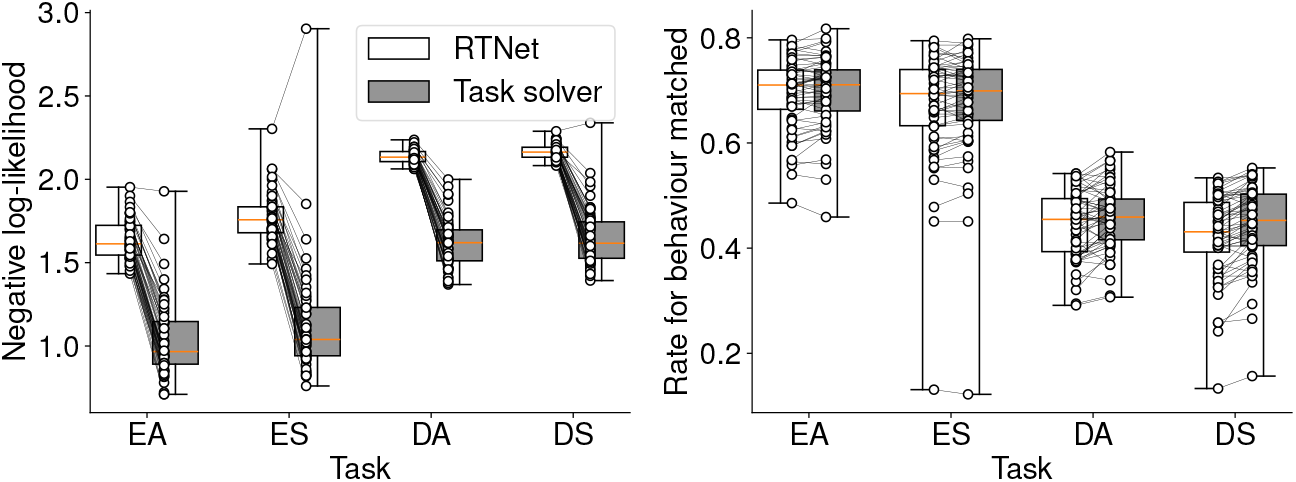
Comparison of prediction performance in Within-Condition Prediction for the MNIST task. The plots show the negative log-likelihood (left) and the rate for behaviour matched (right) for the RTNet model and our task solver. Each connected pair of dots represents a single participant’s data. Box plots indicate the median and interquartile range. Whiskers extend to the minimum and maximum values. The task solver achieves significantly better prediction accuracy.

#### EIDT accurately transfers individuality

Next, in Cross-Condition Transfer, we tested individuality transfer across all 12 pairs of experimental conditions. The full EIDT framework was compared againt a baseline: a task solver (source) model trained directly on a test participant’s source condition data. The EIDT framework consistently and significantly outperformed this baseline across all transfer sets (Figure 9). A two-way (model: task solver/EIDT, transfer direction: 12 sets (see horizontal axis)) RM ANOVA confirmed a significant effect of the model on negative log-likelihood (model: *F*_3,177_ = 2440.373, *p* < 0.001, 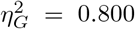, transfer direction: *F*_11,649_ = 347.850, *p* < 0.001, 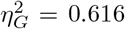, interaction: *F*_33,1947_ = 336.968, *p* < 0.001, 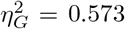) and rate for behaviour matched (model: *F*_3,177_ = 2318.456, *p* < 0.001, 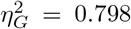, transfer direction: *F*_11,649_ = 394.753, *p* < 0.001, 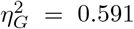, interaction: *F*_33,1947_ = 355.577, *p* < 0.001, 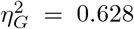). The model was also able to reproduce idiosyncratic error patterns of individual participants, such as Participant #23’s lower accuracy for digit 1 and Participant #56’s difficulty with digits 6 and 7 (Figure 10).

**Figure 9.**
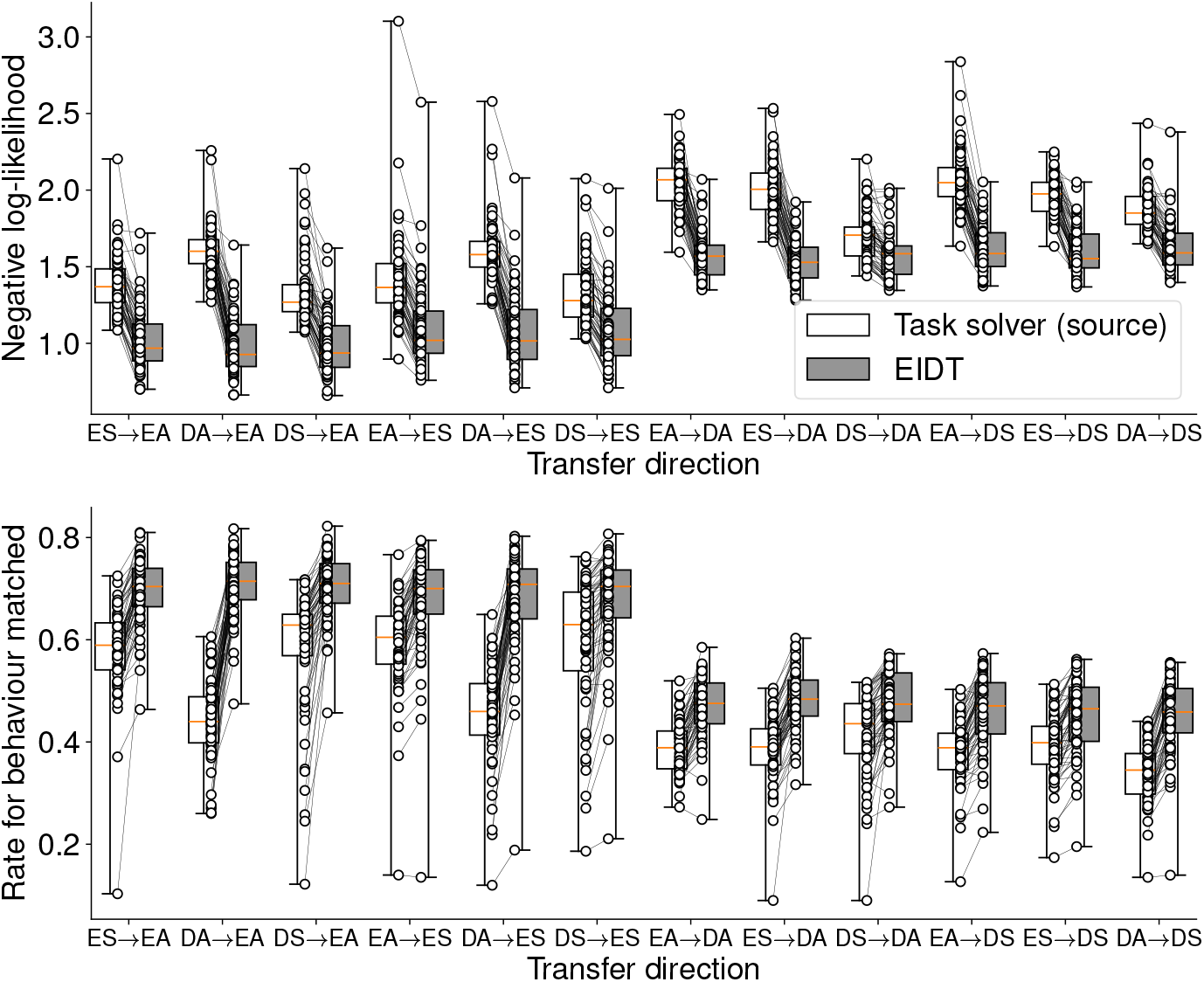
Individuality transfer performance in Cross-Condition Transfer for the MNIST task. The plots compare the EIDT framework against the task solver (source) baseline across all 12 transfer directions on negative log-likelihood (top) and rate for behaviour matched (bottom). Each connected pair of dots represents a single participant’s data. Box plots indicate the median and interquartile range. Whiskers extend to the minimum and maximum values. EIDT consistently demonstrates superior prediction accuracy.

**Figure 10.**
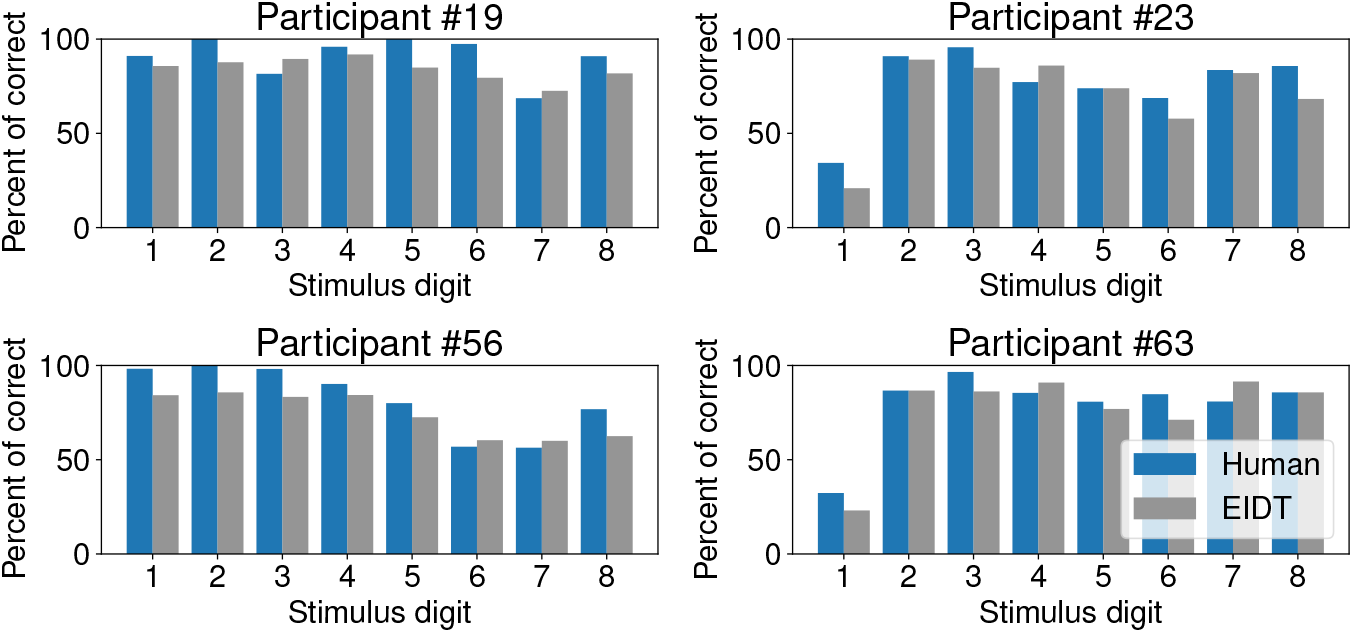
EIDT captures individual-specific error patterns in the MNIST task. The plots show the percentage of correct responses for each digit for four representative participants (blue bars) and their corresponding EIDT-generated models (gray bars). Data shown is for the ES target condition, with transfer from EA.

#### Latent space reflects behavioural tendencies

Similar to the MDP task, a cross-individual analysis showed that the distance in the latent space was significant predictor of prediction performance for all transfer directions (Figure 11; see Figures S8, S9, and Table S2, Supplementary Materials, for full results). This confirms that, in the perceptual domain as well, the individual latent representation captures meaningful behavioural differences that are critical for accurate prediction.

**Figure 11.**
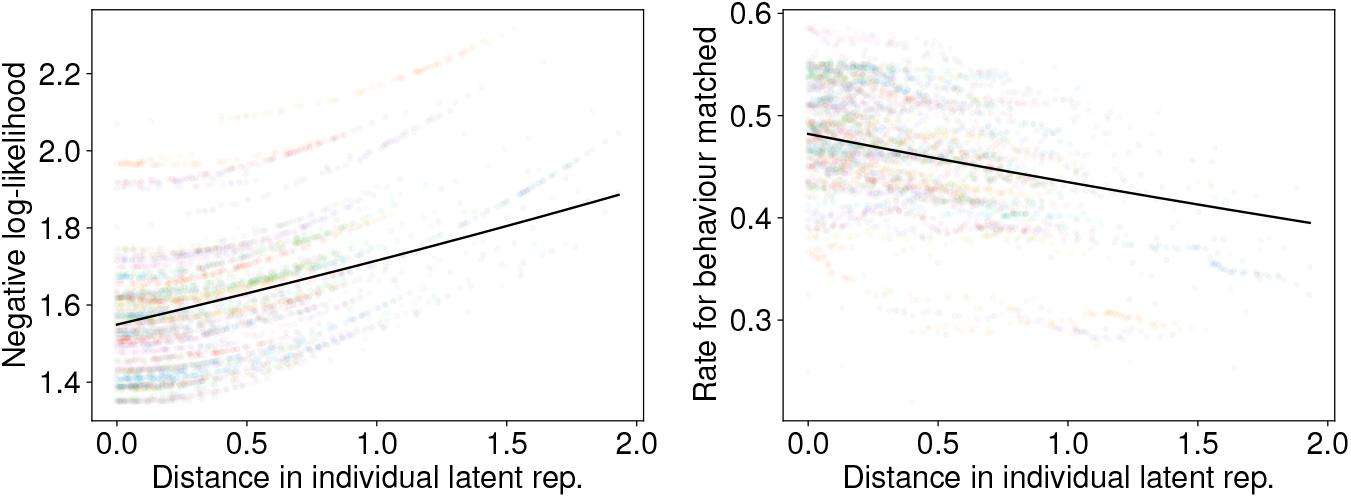
Prediction performance as a function of latent space distance in the MNIST task (transfer direction EA → DA). This cross-individual analysis shows the result of using a task solver generated from one participant to predict the behaviour of another participant. The horizontal axis is the Euclidean distance between the latent representation of the two participants. The vertical axis shows the negative log-likelihood (left) and rate for behaviour matched (right). Each dot represents one participant pair. Performance degrades as the distance between individuals increases, with the solid line showing the GLM fit.

## 3 Discussion

We proposed an EIDT framework for modeling the unique decision-making process of each individual. This framework enables the transfer of an individual latent representation from a (source) task condition to a different (target) task condition, allowing a task solver predict behaviours in the target task condition. Several neural network techniques, such as autoencoders [38, 48], hypernetworks [20], and learning-to-learn [53, 43], facilitate this transfer. Our experiments, conducted on both value-guided sequential and perceptual decision-making tasks, demonstrated the potential of the proposed framework in individuality transfer across task conditions.

### EIDT framework extends prior work on individuality transfer

The core concept of using an encoder-decoder architecture to capture individuality builds on the work of Dezfouli et al. [10], who applied a similar model to a bandit task. We extended this idea in three key ways. First, we validated that the framework is effective for previously unseen individuals who were not included in model training. Although these individuals provided behavioural data in the source task condition to identify their individual latent representations, their data were not used for model training. Second, we established that this transfer is effective across different experimental conditions (e.g., changes in task rules or difficulty), not just across sessions of the same task. Third, while the original work focused on value-guided tasks, we validated the framework’s applicability to perceptual decision-making tasks, specifically the MNIST task. These findings establish that EIDT effectively captures individual differences across both task conditions and individuals.

### Interpreting the individual latent representation remains challenging

Although we found that Q-learning parameters were reflected in the individual latent representation, the interpretation of this representation remains an open question. Since interpretation often requires task-condition-specific considerations [13], it falls outside the primary scope of this study, whose aim is to develop a general framework for individuality transfer. Previous research [28, 18] has explored associating neural network parameters with cognitive or functional meanings. Approaches such as disentangling techniques [2] and cognitive model integration [19, 49, 44, 14] could aid in better understanding the cognitive and functional significance of the individual latent representation.

Regarding the individual latent representation, disentanglement and separation losses [10] during the model training could enhance interpretability. However, we used only the reproduction loss, as defined in (5), because interpretable parameters in cognitive models (e.g., [9]) are not necessarily independent (e.g., an individual with a high learning rate may also have a high inverse temperature [27], resulting these two parameters being represented with one variable).

### Why can the encoder extract individuality for unseen individuals?

Our experiments, which divided participants into training and test participant pools, demonstrated that the framework successfully extracts individuality for completely new individuals. This generalization likely relies on the fact that individuals with similar behavioural patterns result in similar individual latent representation and individuals similar to new participants exist in the training participant pool [57]. This hypothesis suggests that individuals can be clustered based on behavioural patterns. Behavioural clustering has been widely discussed in relation to psychiatric conditions, medication effects, and gender-based differences (e.g., [31, 50, 40]). Our results could contribute to a deeper discussion of behavioural characteristics by clustering not only these groups but also healthy controls.

### Which processes contribute to individuality?

In the MNIST task, we assumed that individuality emerged primarily from the decision-making process (implemented by an RNN [45, 6]), rather than from the visual processing system (implemented by a CNN [55]). The CNN was pretrained, and the decoder did not tune its weights. Our results do not rule out the possibility that the visual system also exhibits individuality [24, 47]; however, they imply that individual differences in perceptual decision-making can be explained primarily by variations in the decision-making system [36, 51, 57, 21]. This assumption provides valuable insights for research on human perception.

### Limitations

One limitation is that the source and target behaviours were performed on different conditions, but within the same task. Thus, our findings do not fully evaluate the generalizability of individuality transfer across diverse task domains. However, our framework has the potential to be applied to diverse tasks since it connects the source and target tasks via the individual latent representation and accepts completely different tasks for the source and target. A key to realizing this transfer might be ensuring that the cognitive functions, such as memory, required for solving the source and target tasks are (partially) shared. The latent representation is expected to represent individual features of these functions. Conversely, if source and target tasks require completely different functions to solve them, the transfer by EIDT would not work.

The effectiveness of individuality transfer may be influenced by dataset volume. As discussed earlier, prediction performance may depend on whether similar individuals exist in the training participant pool. In our study, 100 participants were sufficient for effective transfer. However, tasks involving greater behavioural diversity may require a substantially larger dataset.

As discussed earlier, the interpretability of the individual latent representation requires further investigation. Furthermore, the optimal dimensionality of the individual latent representation remains unclear. This likely depends on the complexity of tasks involved—specifically, the number of factors needed to represent the diversity of behaviour observed in those tasks. While these factors have been explored in cognitive modeling research (e.g., [23, 13]), a clear understanding at the individual level is still lacking. Integrating cognitive modeling with data-driven neural network approaches [10, 19] could help identify key factors underlying individual differences in decision-making.

### Future directions

To further generalize our framework, a large-scale dataset is necessary, as discussed in the limitations. This dataset should include a large number of participants to ensure prediction performance for diverse individuals [32]. All participants should perform the same set of tasks, which should include a variety of tasks [56]. Building upon our framework, where the encoder currently accepts action sequences from only a single task, a more generalizable encoder should be able to process behavioural data from multiple tasks to generate a more robust individual latent representation. To enhance the encoder, a multi-head neural network architecture [4] could be utilized. A individual latent representation would enable transfer to a wider variety of tasks and allow accurate and detailed parameterization of individuals using data from only a single task.

Robust and generalizable parameterization of individuality enables computational modeling at the individual level. This approach, in turn, makes it possible to replicate individuals’ cognitive and functional characteristics *in silico* [41]. We anticipate that it offers a promising pathway toward a new frontier: artificial intelligence endowed with individuality.

## 4 Methods

### 4.1 General framework for individuality transfer across task conditions

We formulate the problem of individuality transfer, which involves extracting an individual latent representation from a source task condition and predicting behaviour in a target task condition with preserving individuality. We consider two task conditions, *A* and *B*, which are different but related. For example, condition *A* might be a 2-step MDP, while condition *B* is a 3-step MDP.

The individuality transfer across task conditions is defined as follows. An individual *K* performs a problem within condition *A*, with their behaviour recorded as 𝒜_*K*_. Our objective is to predict ℬ_*K*_, which represents *K*’s behaviour when performing a task with condition *B*. To achieve this, we extract an individual latent representation ***z*** from 𝒜_*K*_, capturing the individual’s behavioural characteristics. This representation ***z*** is then used to construct a task solver, enabling it to mimic *K*’s behaviour in condition *B*. Since condition *A* provides data for estimating the individual latent representation and condition *B* is the target of behaviour prediction, we refer to them as the source task condition and target task condition, respectively.

Our proposed framework for the individuality transfer consists of three modules:

**Task solver** predicts behaviour in the target condition *B*.

**Encoder** extracts the individual latent representation from the source condition *A*.

**Decoder** generates the weights of the task solver based on the individual latent representation.

These modules are illustrated in Figure 1. We refer to this framework as EIDT, an acronym for encoder, individual latent representation, decoder and task solver.

#### 4.1.1 Data representation

For training, we assume that behaviour data from a participant pool 𝒫 (*K* ∉ 𝒫), where each participant has performed both conditions *A* and *B*. These datasets are represented as 𝒜 = {𝒜_*n*_}_*n*∈*P*_ and ℬ = {*ℬ*_*n*_}_*n*∈*P*_.

For each individual *n*, the set 𝒜_*n*_ consists of one or more sets, each containing a problem instance *ϕ* (stimuli, task settings, or environment in condition *A*) and a sequence of action(s) *α* (recorded behavioural responses). For example, in an MDP task, *ϕ* represents the Markov process (state-action-reward transition) and *α* consists of choices over multiple trials. In a simple object recognition task, *ϕ* is a visual stimulus and *α* is the participant’s response to the stimulus. Similarly, *B*_*n*_ consists of a problem instance *ψ* and an action sequence *β*.

#### 4.1.2 Task solver

The task solver predicts the action sequence for condition *B* as

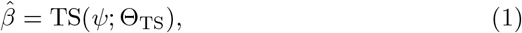

where *ψ* is a specific problem in condition *B* and Θ_TS_ represents the solver’s weights. The task solver architecture is tailored to condition *B*. For example, in an MDP task, the task solver outputs a sequence of actions in response to *ψ*. In a simple object recognition task, it produces an action based on a visual stimulus *ψ*.

#### 4.1.3 Encoder

The encoder processes an action sequence(s) *α* and generates an individual latent representation ***z*** ∈ ℝ^*M*^ as

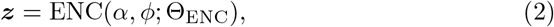

where *ϕ* is a problem in condition *A*, Θ_ENC_ represents the encoder’s weights, and *M* is the dimensionality of the individual latent representation. The encoder architecture is task-condition-specific and designed for condition *A*.

#### 4.1.4 Decoder

The decoder receives the individual latent representation ***z*** and generates the task solver’s weights as

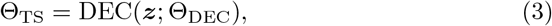

where Θ_DEC_ represents the decoder’s weights. Since the decoder determines the task solver’s weights, it functions as a hypernetwork [20, 22].

#### 4.1.5 Training objective

Although conditions *A* and *B* differ, an individual’s decision-making system remains consistent across task conditions. We model this using the individual latent representation ***z***, linking it to the task solver via the encoder and decoder. For training, we use behavioural dataset {𝒜_*n*_, ℬ_*n*_}_*n*∈*P*_ from a individual pool 𝒫.

Let *α* be an action sequence representing individual *n*’s behaviour on the source task condition, i.e., (*α, ϕ*) ∈ 𝒜_*n*_, *n* ∈ 𝒫. The individual latent representation is derived by ***z*** = ENC(*α, ϕ*; Θ_ENC_). The weights of the task solver are then given by Θ_TS_ = DEC(***z***; Θ_DEC_). Subsequently, the task solver, with the given weights, predicts an action sequence for condition *B* as 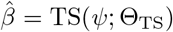, where (*β, ψ*) ∈ ℬ_*n*_. We then measure the prediction error between 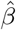 and *β* as:

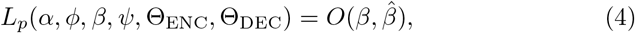

where *β* is an action sequence in ℬ_*n*_ recorded along with the problem *ψ*, and *O*(·,·) is a suitable loss function (e.g., likelihood-based loss for probabilistic outputs). Using the datasets containing the behaviour of the individual pool, 𝒫 the weights of the encoder and decoders, Θ_ENC_ and Θ_DEC_, are optimized by minimizing the total loss:

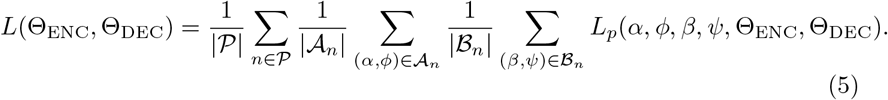

This section provides a general formulation of individuality transfer across two task conditions. For specific details on task architectures and loss functions, see Sections 4.2 and 4.3.

### 4.2 Experiment on MDP task

We validated our individuality transfer framework using two different decision-making tasks: the MDP task and the MNIST task. This section focuses on the MDP tasks, a dynamic multi-step decision-making task.

#### 4.2.1 Task

At the beginning of each episode, an initial state-cue is presented to the participant. For human participants, the state-cue is represented by animal images (Figure 12). For the cognitive model (Q-learning agent) and neural network-based model, the state-cue is represented numerically (e.g., (2, 1) for the first task state in the second choice). The participant makes a binary decision (denoted as action *C*_1_ or *C*_2_) for each step. In the human experiment, these actions correspond to pressing the left or right cursor key. With a certain probability (either 0.8/0.2 or 0.6/0.4), known as the state-action transition probability, the participant transitions to one of two subsequent task states. This process repeats two times for the 2-step MDP and three times in the 3-step MDP. After the final step, the participant receives an outcome: either a reward (*r* = 1) or no reward (*r* = 0). For human participants, rewards were displayed as symbols, as shown in Figure 12. Each sequence from initial state-cue presentation to reward delivery constitutes an *episode*.

**Figure 12.**
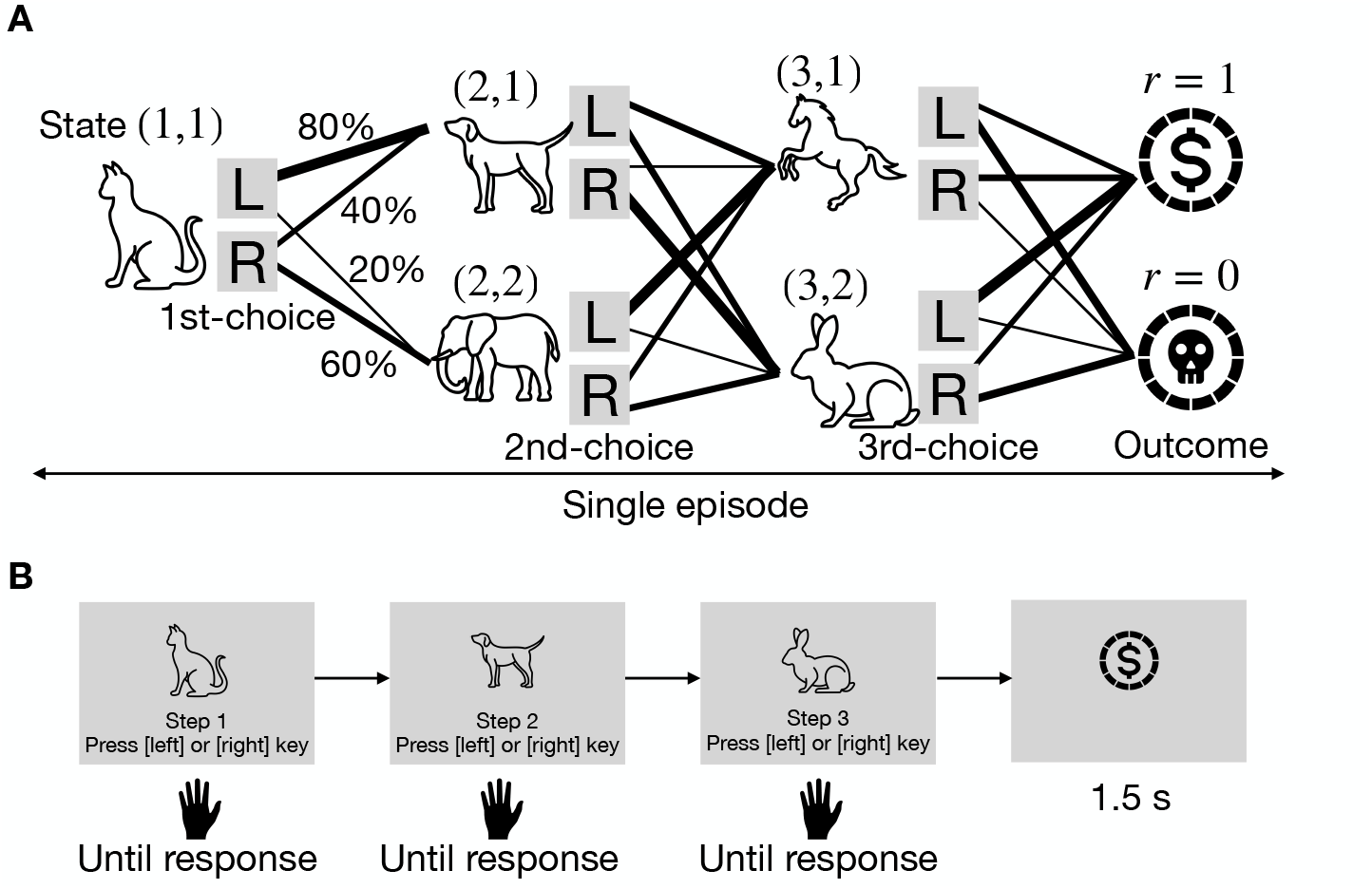
The 3-step MDP task. **A**. Tree diagram illustrating state-action transitions. **B**. Flow of a single episode in the behavioural experiment for human participants.

The state-action transition probability *T* (*s, a, s*^*′*^) from a task state *s* to a preceding state *s*^*′*^ given an action *a* varies gradually across episodes. With probability *p*_trans_, one of the transition probabilities switches to a new set chosen from {(0.8, 0.2), (0.2, 0.8), (0.6, 0.4), (0.4, 0.6)}. Consequently, participants must adjust their decision-making strategy in response to these shifts in transition probabilities to maintain reward maximization.

#### 4.2.2 Behavioural data collection

We recruited 123 participants via Prolific. All participants provided their informed consent online. This study was approved by the Committee for Human Research at the Graduate School of Engineering, The University of Osaka, and compiled with the Declaration of Helsinki. Participants received a base compensation of £4 for completing the entire experiment, A performance-based bonus (£0 to £2, average: £1) was awarded based on rewards earned in the MDP task. Each participant completed 3 sequences for each step condition (2-step and 3-step MDP tasks), with each sequence comprising 50 episodes. The order of the 2-step and 3-step MDP tasks was randomized across sequences. State-cue assignment (animal images) were randomly determined for each sequence. Participants took a mandatory break (≥ 1 minute) between sequences.

To ensure data quality, we applied exclusion criteria based on average reward, action bias, and response time. Thresholds for these metrics were systematically determined using the interquartile range method on statistics from the initial dataset. Participants were removed from the analysis entirely if their data from any single block fell outside these established ranges. This procedure led to the exclusion of one participant for low average reward (below 0.387 for the 2-step MDP and 0.382 for the 3-step MDP), 23 participants for excessive action bias (outside the 26.3–73.3% range), and 18 for outlier response times (outside the 0.260–1.983 s range). In total, 42 participants (approximately 34%) were excluded, resulting in a final sample of 81 participants for analysis.

#### 4.2.3 Cognitive model: Q-learning

To model decision-making in the MDP task, we employed a Q-learning agent [46]. At each step *t*, the agent was presented with the current task state *s*_*t*_ and selected an action *a*_*t*_. The agent maintained *Q*-values, denoted as *Q*(*s, a*), for all state-action pairs, where *s* was a state of the set of all possible task states *S* and *a* was an action of the set of available actions in that state 𝒞 _*s*_. The probability of selecting action *a* was determined by a softmax policy:

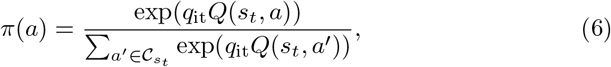

where *q*_it_ > 0 was a parameter called the inverse temperature or reward sensitivity, controlling the balance between exploration and exploitation.

After selecting action *a*_*t*_, the agent received an outcome *r*_*t*_ ∈ {0, 1} and transitioned to a new state *s*_*t*+1_. The *Q*-value for the selected action was updated by

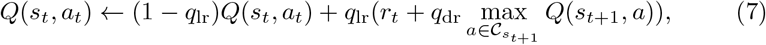

where *q*_lr_ ∈ (0, 1) was the learning rate, determining how much newly acquired information replaced existing knowledge, and *q*_dr_ ∈ (0, 1) was the discount rate, governing the extent to which future rewards influenced current decision. The *Q*-values are initialized as *q*_init_ before an agent starts first episode.

#### 4.2.4 EIDT model

This section describes the specific models used for individuality transfer in the MDP task.

##### Data representation

Since MDP tasks involve sequential decision-making, each action sequence consists of multiple actions within a single session. In our experiment, each participant completed *L* trials per session, with *L* = 100 for the 2-step MDP and *L* = 150 for the 3-step MDP. The action sequence is represented as [(*s*_1_, *a*_1_, *r*_1_), …, (*s*_*L*_, *a*_*L*_, *r*_*L*_)], where, *s*_*t*_ denotes the task state at trial *t, a*_*t*_ ∈ *C* represents the action selected from the set 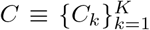 (with *K* = 2 in our task), and *r*_*t*_ ∈ {0, 1} indicates whether a reward was received. In the *M*-step MDP described in Section 4.2, each task state is represented as (*m, c*_*m*_), where *m* denotes the current step within the episode (*m* ∈ {1, …, *M*}) and *c*_*m*_ corresponds to the cue presented to the participant. The action sequence, denoted as *α* or *β*, consists of a sequence of selected actions (*a*_1_, …, *a*_*L*_), while a problem, denoted as *ϕ* or *ψ*, is represented as ((*s*_1_, …, *s*_*L*_), (*r*_1_, …, *r*_*L*_)).

##### Task solver

Before describing the encoder and decoder, we define the architecture of the task solver, which generates actions for the *M*-step MDP task. The task solver is implemented using a gated recurrent unit (GRU) [7] with *Q* cells, where *Q* = 4 for the 2-step task and *Q* = 8 for the 3-step task. At time-step *t*, the GRU takes as input the previous hidden state ***h***_*t*−1_ ∈ ℝ^*Q*^, the previous task state *s*_*t*−1_, the previous action *a*_*t*−1_, the previous reward *r*_*t*−1_, and the current task state *s*_*t*_. It then updates the hidden state as

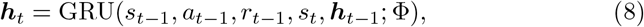

where Φ represents the GRU’s weights. The updated hidden state is then used to predict the probability of selecting each action through a fully-connected feed forward layer:

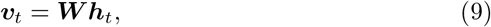

where ***v***_*t*_ represents the logit scores for each action (unnormalised probabilities), and ***W* ∈** ℝ^*K×Q*^ is the weight matrix. The probabilities of each action are computed using a softmax layer:

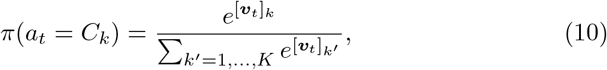

where *π*(*a*_*t*_ = *C*_*k*_) represents the probability of selecting action *C*_*k*_ at time *t*, and [***v***_*t*_]_*i*_ denotes the *i*-th element of ***v***_*t*_.

For input encoding, we used a 1-of-K scheme. The step of the MDP task is encoded as [1, 0, 0] for step 1, [0, 1, 0] for step 2, and [0, 0, 1] for step 3. Each task state *s*_*m*_ is represented as [1, 0] or [0, 1] to distinguish the two state cues at each step. The participant’s action is encoded as *C*_1_: [1, 0] or *C*_2_: [0, 1], while the reward is represented as 0: [1, 0] or 1: [0, 1]. These encodings are concatenated to form input sequences.

The task solver TS(*ψ*; Θ_TS_) generates a sequence of predicted action probabilities 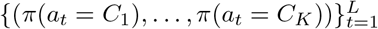, using the GRU, the fully-connected layer ***W***, and the softmax layer. The problem *ψ* defines the MDP environment, specifying state transitions and reward outcomes in response to selected action. To evaluate prediction accuracy, the loss function *O*(*β*,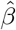), defined in (4), compares human-performed action {*β, ψ*} with those predicted by the task solver, 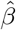, *ψ*. Notably, the problem *ψ* is not executed with the task solver; instead, the task solver predicts action probabilities based on the same task state and reward history as in the human behavioural data.

##### Encoder and decoder

The encoder ENC(*α, ϕ*; Θ_ENC_) extracts an individual latent representation ***z*** from a sequence of actions *α* corresponding to a given environment *ϕ*. The first module of the encoder is a GRU, similar to the task solver, with *R* = 32 cells. The final hidden state ***h***_*L*_ ∈ ℝ^*R*^ serves as the basis for computing the individual latent representation [***z*** ∈ ℝ^*M*^ using a fully-connected feed-forward network with four layers *d*(·) as ***z*** = *d*(***h***_*L*_).

The decoder takes the individual latent representation ***z*** as input and generates the weights for the task solver by Θ_TS_ = DEC(***z***; Θ_DEC_). The decoder is implemented as a single-layer linear network.

### 4.3 Experiment on MNIST task

This section describes the specific models used for individuality transfer in hand-written digit recognition (MNIST) task.

#### 4.3.1 Task

The dataset used in this experiment was originally collected and published by Rafiei et al. [34]. In this task, participants were presented with a stimulus image depicting a handwritten digit and were required to respond by pressing the corresponding number key, as illustrated Figure 1. For further details regarding the task design and data collection, refer to [34].

#### 4.3.2 EIDT model

##### Data representation

An action sequence, denoted as *α* or *β*, consists of a single action *a* and its corresponding response time *b*. The associated problem, represented as *ϕ* or *ψ*, corresponds to a stimulus image. The action *a* is selected from a set {*C*_1_, … *C*_*K*_}. Since the task involves recognizing digits ranging from 0 to 9, the number of possible actions is *K* = 10. The stimulus image, *ϕ* or *ψ*, is an image of size *H* × *W*. In this experiment, we adopted the same resolution as [34], setting *H* = *W* = 227.

##### Task solver

The task solver for the handwritten digit recognition task is based on the model proposed by [34]. Their model consists of a CNN and an evidence accumulation module. However, since their model represents average human behaviour and does not account for individuality differences, we replace the accumulation module with a GRU [6] to capture individuality. The CNN module processes the input image and produces an evidence vector ***e*** = CNN(*ψ*), where ***e*** ∈ ℝ^*K*^ and CNN(·) is based on the AlexNet architecture [26]. The weights of the CNN are sampled from a Bayesian neural network (BNN), introducing stochasticity in the output. This stochasticity enables the models to generate human-like, probabilistic decisions.

The stimulus image is fed into the CNN *S* times, generating *S* evidence distributions ***e***_*t*_ ∈ ℝ^*K*^ at each time step *t* = 0, …, *S* −1. In this study, we set *S* = 16 to match the maximum response time, as described later. Since the CNN weights are stochastically sampled from the BNN, the CNN’s output varies even when the same image is input multiple times. To model individuality in decision-making, we introduce a GRU with *Q* cells (*Q* = 4 in our setup). The GRU receives as input the previous hidden state ***h***_*t*−1_ ∈ ℝ^*Q*^ and the current evidence ***e***_*t*_, updating its hidden state as

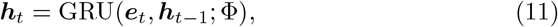

where Φ represents the GRU’s network weights. The updated hidden state is passed through a dense layer (as defined in (9)) and a softmax layer (as defined in (10)) to generate the probability distribution over possible digit classifications [*P*_*t*_(*C*_1_), …, *P*_*t*_(*C*_*K*_)] at each time step *t*.

To evaluate the prediction error, we compare the action sequences generated by human participants {*β, ψ*}with those predicted by the task solver {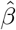, *ψ* }, incorporating response times into these analysis. The actual response time *b* is converted into an integer time step 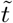 using the formula: 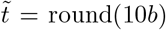. For example, a response time of *b* = 0.765 s is converted to 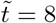. The likelihood of observed decision is then calculated as 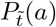, where *a* is the actual digit chosen by the participant.

In this task solver, the CNN (driven by BNN) models a visual processing system, while the RNN represents the decision-making system. We assume that the visual system (implemented by CNN and BNN) is shared across all individuals, whereas the decision-making system (implemented by RNN) captures individual differences. Based on this assumption, the CNN and BNN are pretrained using the MNIST dataset [26], and their weight distributions are fixed across individuals. The pretraining procedure followed the original methodology [34].

##### Encoder and decoder

Since each action sequence contains only a single action, it does not form a true “sequence.” This makes it challenging to extract individuality from a single data point. To address this, the encoder takes a set of single action sequences as input rather than a single sequence. Specifically, the encoder 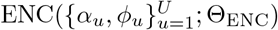 extracts the individual latent representation ***z*** from *U* sets of stimulus images *ϕ*_*u*_ and their corresponding responses *α*_*u*_, where *u* = 1, …, *U*. Here, *ϕ*_*u*_ represent the stimulus presented in the *u*-th trial, and *α*_*u*_ represents the corresponding response. The number of action sequences *U* corresponds to the number of samples available for each individual in the dataset. Since the outputs for these action sequences are just averaged, *U* can be adjusted flexibly.

The encoder architecture consists of a single CNN module, a single GRU, and a fully-connected feed-forward network. The CNN module is identical to the one used in the task solver. Given an input *ϕ*_*u*_, let ***e***_*t,u*_ represent the evidence output from the CNN at time step *t*. The GRU, which consists *R* cells (*R* = 16 in our setup), updates its hidden state based on the previous state, the current CNN evidence, and an encoding of the response action by

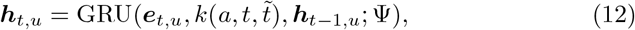

where Ψ represents the network weights. The function *k*(*a, t*, 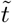) outputs the one-hot encoded action *a* if 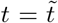, and zeros otherwise. The value 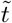 represents the converted response time, obtained from the original response time *b* in the action sequence *α*_*u*_. After processing all *U* sequences, the final hidden states are averaged across sequences: 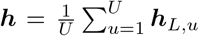. The individual latent representation is then computed as ***z*** = *d*(***h***), where *d*(·) represents a single-layer fully-connected feed-forward network. The decoder, implemented as a single linear layer, takes the individual latent representation ***z*** as input and outputs the weights for the task solver.

## 5 Data availability

The behavioural data for the MDP task has been made publicly available at https://github.com/hgshrs/indiv_trans

## 6 Code availability

All code and trained models have been made publicly available at https://github.com/hgshrs/indiv_trans

## Acknowledgements

This work was supported in part by the Japan Society for the Promotion of Science (JSPS) KAKENHI, grant number 22H05163 and 24K15047, and Japan Science and Technology Agency (JST) Advanced International Collaborative Research Program (AdCORP), grant number JPMJKB2307. We appreciate Kaede Hashiguchi and Yuichi Tanaka, Graduate School of Engineering, The University of Osaka, who gave useful comments for this research.

## Author contributions

H.H. designed and performed the research, collected a part of the data, analyzed the data, and drafted and edited the paper.

