## Supplemental materials for "Individuality transfer: Predicting human decision-making across task conditions"

### Supplementary materials for "Individuality transfer: Predicting human decision-making across task conditions"

Hiroshi Higashi[1]

[1]Graduate School of Engineering, The University of Osaka

October 23, 2025

#### S1 Additional results on MDP task

**Parameter fitting of Q-learning for human behaviours** Figure S1 shows the parameters for the Q-learning model estimated from the behavioural data of human participants. The parameters were estimated separately for the 2-step and 3-step tasks. While the learning rate ($q_{\mathrm{lr}}$) and inverse temperature ($q_{\mathrm{it}}$) were distributed across a range of values, the discount rate ($q_{\mathrm{dr}}$) and the initial Q-value $q_{\mathrm{init}}$ were concentrated near 1 and 0, respectively.

**EIDT training** The losses for the training and validation samples during the EIDT network training are shown in Figure S2. Since we adopted a leave-one-participant-out cross-validation, which results in many training runs, the displayed curves are representatives from a training run that used all participants as training data to illustrate the general convergence pattern. Training was stopped when the validation loss reached its minimum.

**Individual latent representation** Figure S3 visualizes the individual latent representations computed from the behaviours of human participants (squares) and simulated Q-learning agents (dots). Because a different encoder was trained for each fold of the leave-one-participant-out cross-validation, the displayed representations are from a model trained on all participants' data for illustrative purposes.

**Analysis of individual latent representation with a cognitive model** To interpret the latent space, we analyzed the relationship between the Q-learning parameters of simulated agents and their corresponding individual latent representations. We fitted each dimension of the latent representation ($z_i$)

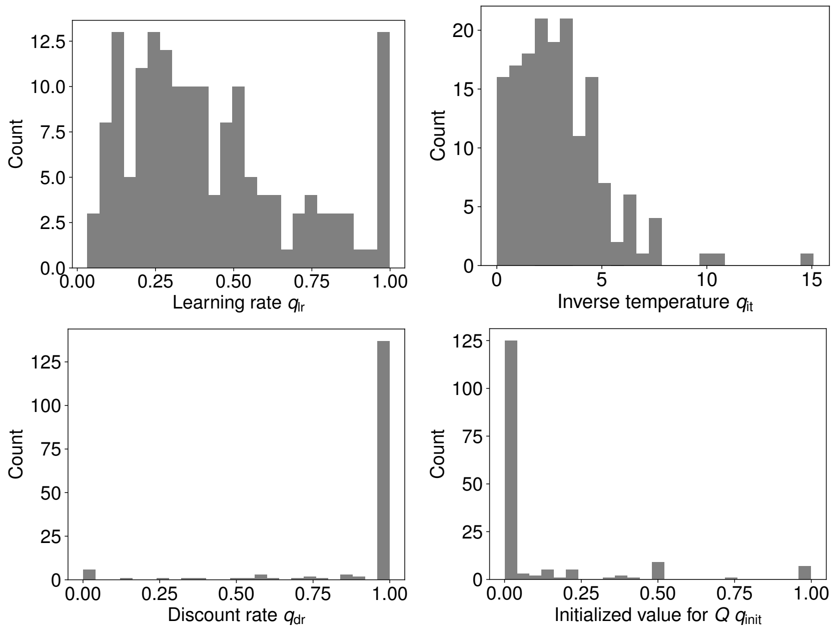


Figure S1: Histograms of Q-learning parameters estimated from human participants' behaviours in the MDP tasks. The distributions for the learning rate and inverse temperature show considerable inter-individual variability, whereas the discount rate and initial Q-value are relatively consistent across participants.

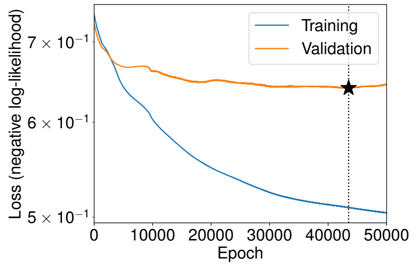


(a) 3-steps to 2-step transfer

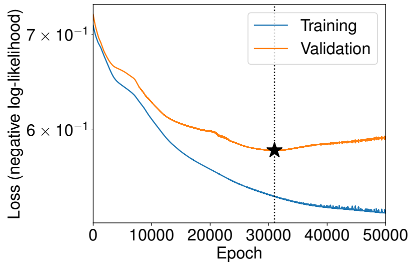


(b) 2-steps to 3-step transfer

Figure S2: Representative training and validation curves for the EIDT model in the MDP task. The plots show the negative log-likelihood loss over training epochs for (a) 3-step to 2-step transfer and (b) 2-step to 3-step transfer. The star marker indicates the point of early stopping, where the validation loss was minimal.

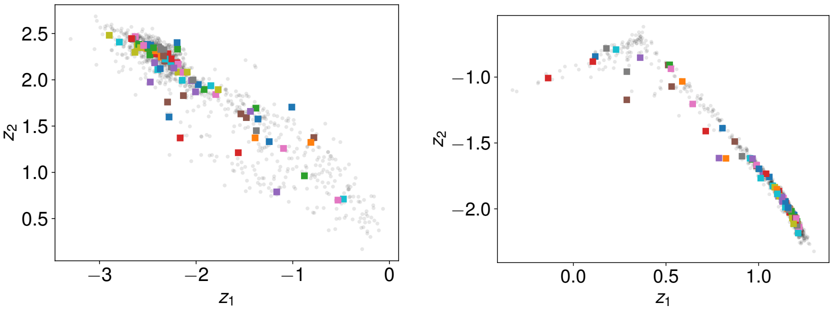


(a) 3-steps to 2-step transfer

(b) 2-steps to 3-step transfer

Figure S3: Individual latent representations for the MDP task. The plots show the two-dimensional latent space derived from behaviours in (a) the 3-step task and (b) the 2-step task. Square markers represent human participants, and dot markers represent simulated Q-learning agents.

Table S1: GLM fitting coefficients for the relationship between Q-learning parameters and the individual latent representation. An asterisk (*) denotes statistical significance ($p < 0.05$).

| | | Coefficients | | | |
|---|---|---|---|---|---|
| Source | Variable | $\beta_0$ (bias) | $\beta_1$ ($q_{\text{lr}}$) | $\beta_2$ ($q_{\text{it}}$) | $\beta_3$ ($q_{\text{lr}} \times q_{\text{it}}$) |
| 2-step | $z_1$ | −0.613* | 0.096* | 0.064* | −0.035* |
| | $z_2$ | 0.634* | −0.079* | −0.066* | 0.028 |
| 3-step | $z_1$ | 1.157* | −0.388* | −0.121* | −0.148* |
| | $z_2$ | −0.646* | 0.183* | 0.061* | −0.071* |

using a generalized linear model (GLM) with the agents' learning rate ($q_{\text{lr}}$) and inverse temperature ($q_{\text{it}}$) as predictors:

$$z_i \sim \text{Normal}\left(\beta_0 + \beta_1 \log(q_{\text{rl}}) + \beta_2 \log(q_{\text{it}}) + \beta_3 \log(q_{\text{lr}}) \log(q_{\text{it}})\right), \qquad (1)$$

The fitted coefficients are summarized in Table S1, and the mapping for the 2-step MDP task is visualized in Figure S4. This analysis complements Figure 6 in the main text, which shows the same mapping for the 3-step task.

**Relationship between prediction performance and latent space for Q-learning agents** We evaluated how the individual latent representation influenced prediction performance for the simulated Q-learning agents. Similar to the analysis on human data, Figure S5 illustrates the prediction performance as a function of the distance in the individual latent representation space in a cross-individual scenario. Using a GLM, we found that the distance was a significant predictor of both negative log-likelihood (transfer direction 3→2:

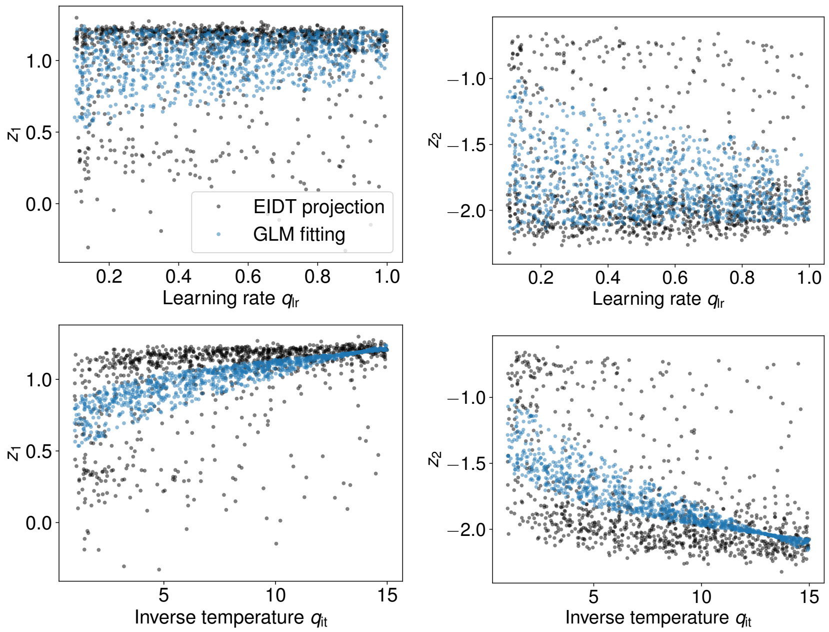


Figure S4: Mapping of Q-learning parameters to the individual latent space for the 2-step MDP task. Each plot shows one dimension of the latent representation ($z_1$ or $z_2$) as a function of either the learning rate ($q_{\mathrm{lr}}$, left) or the inverse temperature ($q_{\mathrm{it}}$, right) of simulated Q-learning agents. Black dots represent the latent representation from the agent's behaviour, while blue dots show the GLM fit.

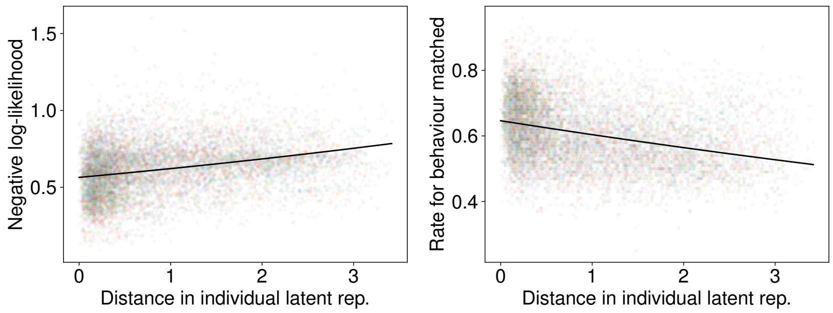


(a) 3-steps to 2-step transfer

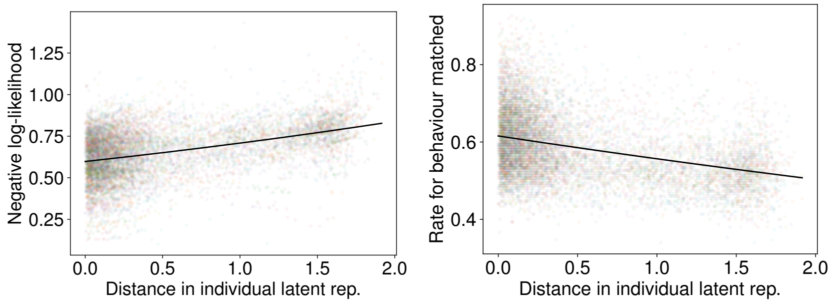


(b) 2-steps to 3-step transfer

Figure S5: Prediction performances for Q-learning agents as a function of latent space distance. The plots show negative log-likelihood (left) and rate for behaviour matched (right) in a cross-individual scenario. (a) 3-step to 2-step transfer. (b) 2-step to 3-step transfer.

$\beta_d = 0.094$, $p < 0.001$, 2→3: $\beta_d = 0.164$, $p < 0.001$) and the rate for behaviour matched (3→2: $\beta_d = -0.069$, $p < 0.001$, 2→3: $\beta_d = -0.097$, $p < 0.001$). This result shows that, as with human data, prediction performance for an agent degrades as the latent distance to the source agent increases, confirming that the latent space captures the behavioural tendencies of the simulated agents.

#### S2 Additional results on MNIST task

**EIDT training** Figure S6 shows representative training and validation loss curves for the EIDT models in the MNIST task. As with the MDP task, these curves are from a model trained from all participants' data for illustrative purposes, showing the typical convergence behaviour.

**Individual latent representation** Figure S7 shows the individual latent representations computed from participants' behaviours for each of the 12 transfer directions in the MNIST task. As we adopted a leave-one-participant-out cross-validation, there were several encoders for each training run. The displayed representations are representatives from a training run using all participants' data.

**Relationship between prediction performance and individual latent representation** We performed a cross-individual analysis for the MNIST task, identical to the one conducted for the MDP task. The prediction performance of a task solver derived from one participant (Participant $l$) was evaluated on the data of another participant (Participant $k$), and this performance was analyzed as a function of the distance between their latent representations ($d_{k,l}$).

The results are shown in Figure S8 (for negative log-likelihood) and Figure S9 (for rate for behaviour matched). In all 12 transfer directions, prediction performance degraded significantly as the distance in the latent space increased. This was confirmed by fitting a GLM:

$$y_{k,l} \sim \text{Gamma}\left(\log(\beta_{\text{participant}_k} + \beta_d d_{k,l} + \beta_0)\right) \tag{2}$$

The fitted coefficients for the distance term ($\beta_d$), shown in Table S2, were significant for all transfer directions ($p < 0.001$), reinforcing the conclusion that the latent space captures meaningful individual differences.

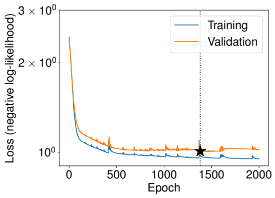


(a) ES→EA

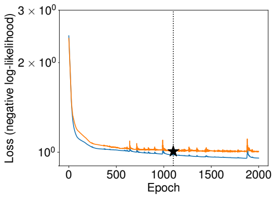


(b) DA→EA

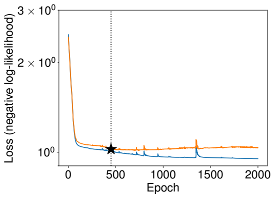


(c) DS→EA

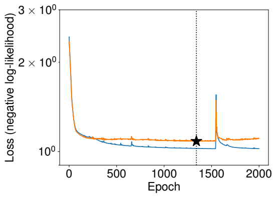


(d) EA→ES

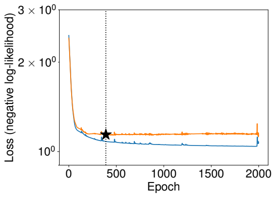


(e) DA→ES

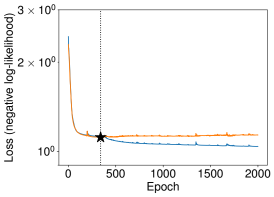


(f) DS→ES

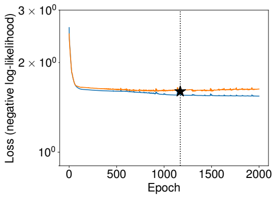


(g) EA→DA

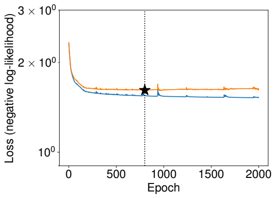


(h) ES→DA

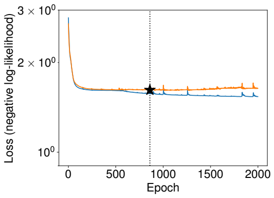


(i) DS→DA

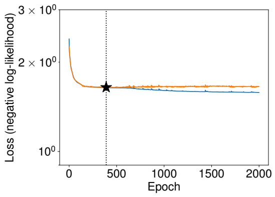


(j) EA→DS

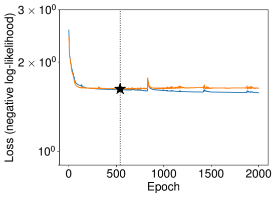


(k) ES→DS

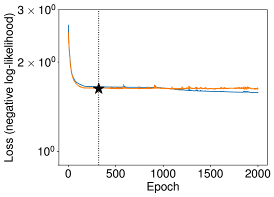


(l) DA→DS

Figure S6: Representatives training and validation curves for EIDT models in the MNIST task for each of the 12 transfer directions. Training was stopped at the epoch with the minimum validation loss, indicated by the start marker.

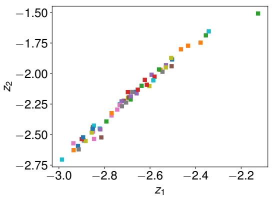

(a) ES→EA

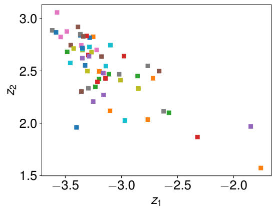

(b) DA→EA

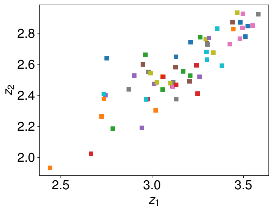

(c) DS→EA

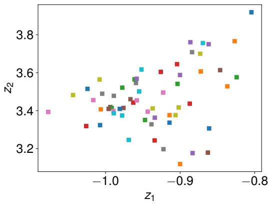

(d) EA→ES

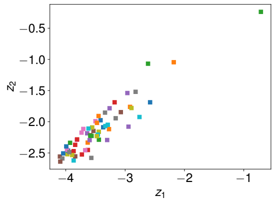

(e) DA→ES

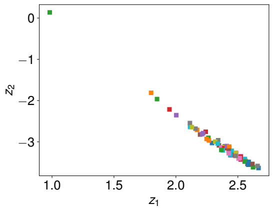

(f) DS→ES

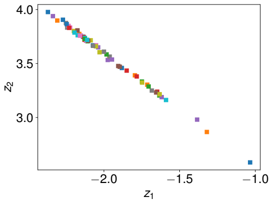

(g) EA→DA

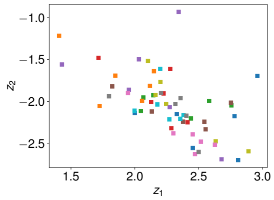

(h) ES→DA

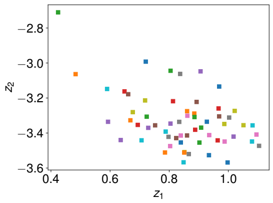

(i) DS→DA

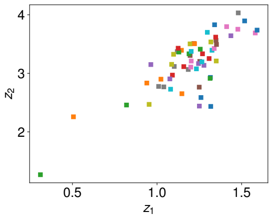

(j) EA→DS

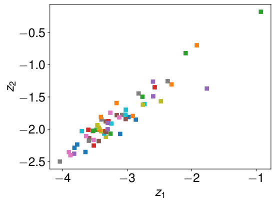

(k) ES→DS

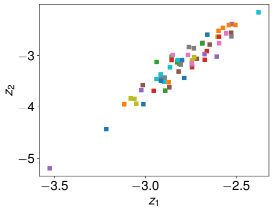

(l) DA→DS

Figure S7: Individual latent representations derived from human participants' behaviours in the MNIST task. Each panel shows the two-dimensional latent space generated when using a different experimental condition as the source. For example, panel (a) shows the latent space when using data from the ES condition to predict behaviour in the EA condition.

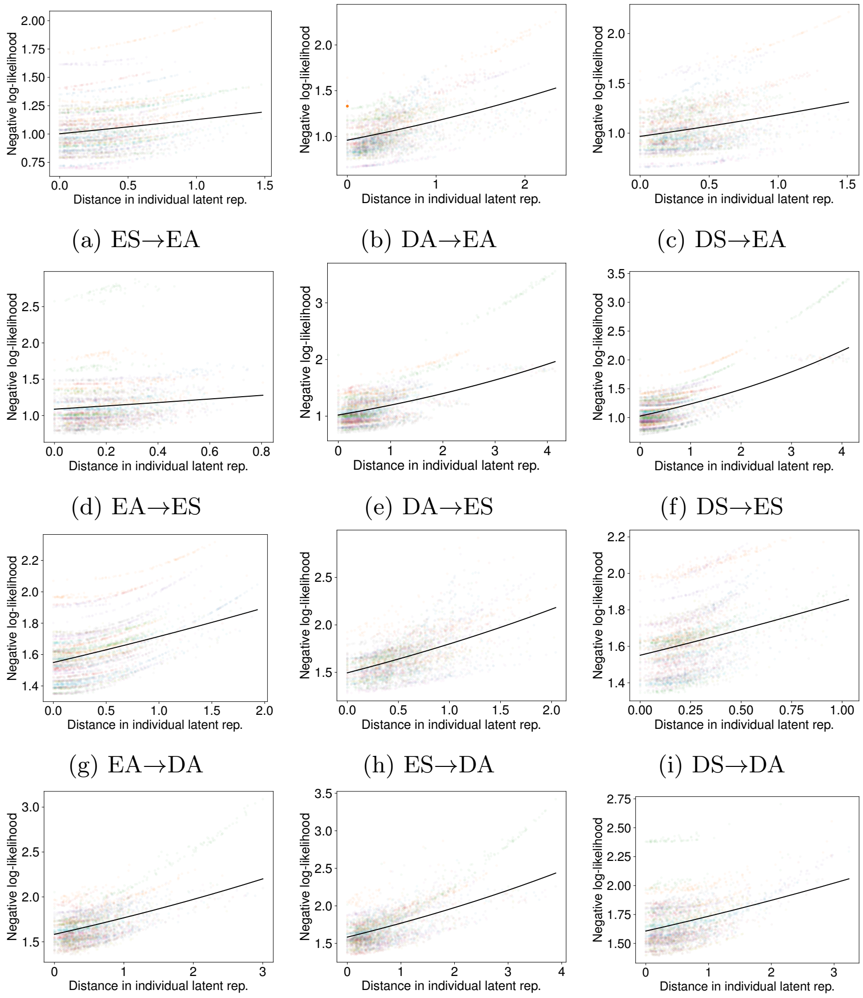


(a) ES→EA (b) DA→EA (c) DS→EA

(d) EA→ES (e) DA→ES (f) DS→ES

(g) EA→DA (h) ES→DA (i) DS→DA

(j) EA→DS (k) ES→DS (l) DA→DS

Figure S8: Prediction performance (negative log-likelihood) as a function of latent space distance in the MNIST task. Each panel shows the results for one of the 12 transfer directions. The negative log-likelihood (vertical axis) increases as the distance between the source and target individuals' latent representations (horizontal axis) increases, indicating worse prediction performance. The solid line is the GLM fit.

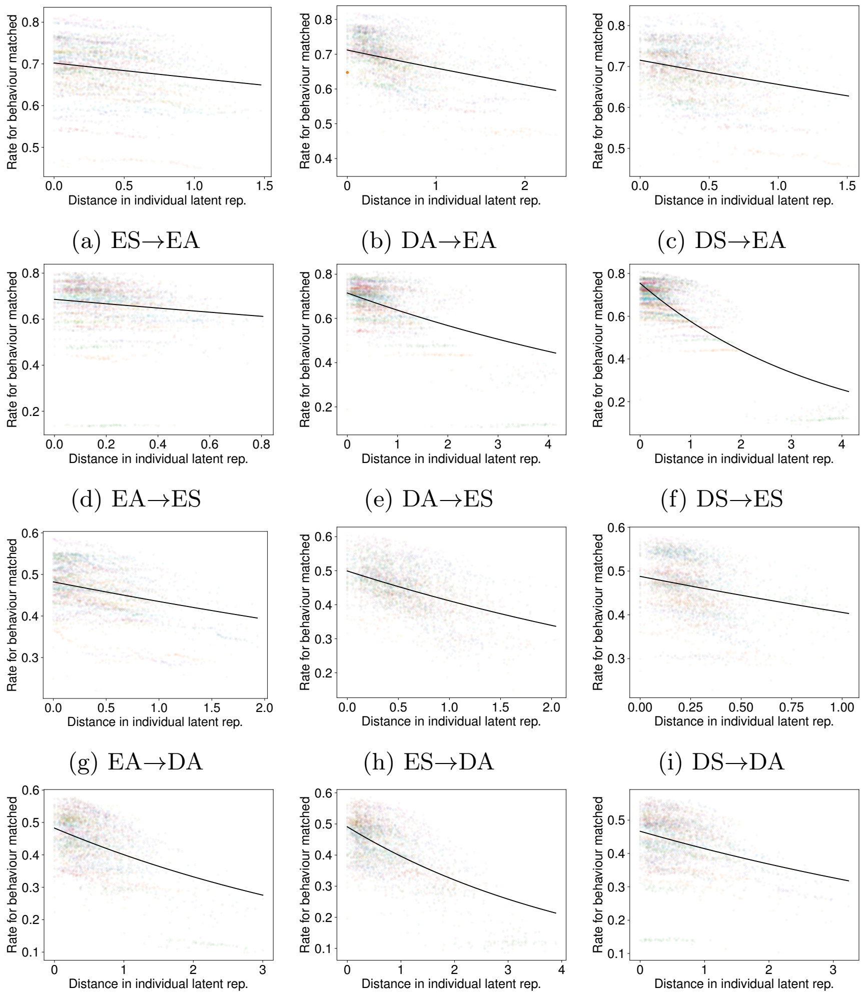


(a) ES→EA (b) DA→EA (c) DS→EA

(d) EA→ES (e) DA→ES (f) DS→ES

(g) EA→DA (h) ES→DA (i) DS→DA

(j) EA→DS (k) ES→DS (l) DA→DS

Figure S9: Prediction performance (rate for behaviour matched) as a function of latent space distance in the MNIST task. Each panel shows the results for one of the 12 transfer directions. The rate for behaviour matched (vertical axis) decreases as the distance between individuals' latent representations (horizontal axis) increases. The solid line is the GLM fit.

Table S2: GLM fitting coefficients ($\beta_d$) for the effect of latent space distance on prediction performance in the MNIST task. All coefficients are statistically significant ($p < 0.001$).

Negative log-likelihood

| | | Target | | | |
|---|---|---|---|---|---|
| | | EA | ES | DA | DS |
| Source | EA | — | 0.202 | 0.102 | 0.110 |
| | ES | 0.117 | — | 0.185 | 0.111 |
| | DA | 0.199 | 0.158 | — | 0.076 |
| | DS | 0.201 | 0.186 | 0.174 | — |

Rate for behaviour matched

| | | Target | | | |
|---|---|---|---|---|---|
| | | EA | ES | DA | DS |
| Source | EA | — | −0.142 | −0.103 | −0.187 |
| | ES | −0.053 | — | −0.193 | −0.214 |
| | DA | −0.076 | −0.115 | — | −0.118 |
| | DS | −0.086 | −0.270 | −0.185 | — |
